# Structural insights into the organization and channel properties of human Pannexin isoforms 1 and 3

**DOI:** 10.1101/2022.09.09.507385

**Authors:** Nazia Hussain, Ashish Apotikar, Shabareesh Pidathala, Sourajit Mukherjee, Ananth Prasad Burada, Sujit Kumar Sikdar, Vinothkumar R. Kutti, Aravind Penmatsa

**Affiliations:** Molecular Biophysics Unit, Indian Institute of Science, Bangalore, 560012; National Centre for Biological Sciences, Tata Institute of Fundamental Research, Bangalore, 560065

**Keywords:** Pannexin1, Pannexin3, ATP-binding, R217H, channel pore, Cryo-EM

## Abstract

Pannexins are single-membrane large-pore ion channels that release ATP upon activation. Three isoforms of pannexins, 1, 2, and 3, perform diverse cellular roles, including inflammation, differentiation, neuropathic pain, and ATP release. In this study, we report the cryoEM structure of pannexin 3 at 3.9 Å and characterize the structural differences with pannexin isoforms 1 and 2. We observe the organization of the Pannexin 3 vestibule into two distinct chambers with a wider pore radius in comparison to both PANX1 and 2 isoforms. We further report the structure of pannexin1 congenital mutant R217H in the resolution range of 3.9 Å. The congenital mutant R217H in transmembrane helix3 (TM3), R217H induce structural changes that leads to a partially closed pore and altered ATP interaction propensities. The channel conductance of the congenital mutant displays weakened voltage sensitivity. The results showcase a complete comparison of the three pannexin isoform structures that along with the structure of Pannexin 1 congenital mutant highlight distinct structural features of pannexin isoforms and the allosteric role of distant substitutions in dictating channel behavior in Pannexin 1.

## Introduction

Pannexins are large-pore vertebrate ion channels identified through sequence similarity with invertebrate gap-junction channels, innexins^1^. Pannexin protomers retain structural and topological similarity to innexins, connexins, and volume-regulated anion channels but remain as single-membrane channels and participate in ion-conductance and ATP release^2^. Three isoforms of pannexins (PANX1, 2 & 3) are identified in mammalian cell types, and each is observed in diverse tissue and physiological niches^3^.

Pannexin1(PANX1) is the more widely studied member among pannexins and is ubiquitously present in several tissue types, and is a channel involved in purinergic signaling through ATP release^4^. It is documented to play a role in diverse processes, including inflammation, cell migration, apoptosis, cytokine secretion, and viral replication^5-8^. The C-terminus of PANX1 is susceptible to caspase 3 cleavage that results in channel-opening and ATP release as a “find-me” signal for macrophages to target apoptotic cells^9^. Besides ATP release, PANX1 is also known to display ionic conductance in response to extracellular potassium, positive potentials, and in some instances display mechanosensitive properties^10,11^. PANX1 knockout results in diverse phenotypes like hearing loss, weakened epileptic propensity, and reduced ATP/cytokine release^12-14^. A recent identification of a congenital mutant of *PANX1* (R217H) that leads to defective channel activity resulting in intellectual disability, ovarian failure, hearing loss, and skeletal defects^15^ signifies the importance of PANX1 as an essential player in human physiology.

A lot less is known about the two other isoforms of PANX1, namely PANX2 and PANX3, which have tissue-specific expression patterns. PANX2 is primarily expressed in neurons and glial cells and is involved in cellular differentiation^16^. PANX3 is expressed in osteoblasts, chondrocytes, and skin and plays a major role in calcium homeostasis suggesting an independent functional niche^17^. Despite the high sequence conservation with its isoform, PANX3 displays distinct structural features, localization, and functional characteristics compared to PANX1_WT_^18^. PANX3 has been observed to regulate osteoblast differentiation through its ER calcium channel function^19^. Along with its predominant role in osteoblast differentiation and wound healing in mice^20^, PANX3 also functions as an ATP release channel similar to PANX1, and the released ATP acts as an activator for P2R-PI3K-AKT signaling^19^. PANX3 has a sequence identity of 42% with PANX1 and is the shortest among the three isoforms with a length of 392 residues^3^. Despite the differences PANX3, is suggested to play a compensatory role upon creating PANX1 knock outs^21^. Multiple attempts to observe PANX3 channel activity in different cellular contexts have been unsuccessful in this channel^22,23^. PANX2 is the most divergent among the three, with a sequence identity of 27% with PANX1, and is characterized by a substantially longer C-terminus^3^. The PANX isoforms, PANX1, PANX2, and PANX3 harbour substitutions in the residues that control channel gating and can have architectural differences that form the basis for altered properties among the three isoforms.

Recent structural studies on different orthologues of Pannexin1 and 2 (PANX1, PANX2) have shown that the channel is organized as a heptamer^24-27^. The topology is similar to large-pore ion channels such as calcium homeostasis modulators (CALHMs), leucine rich repeat containing VRAC subunit 8 (LRRC8), connexins and innexins with four transmembrane helices (TMs), a prominent extracellular domain (ECD), and a disordered C-terminus that is not visible in most PANX1 structures^28^. The access to the channel vestibule for the C-terminus truncated PANX1 and 2 isoforms through the cytosol is unhindered except for a constriction point at the ECD that would enforce selective ion/ATP gating depending on the pore radius. This pore, lined by tryptophan in PANX1 and arginine in PANX2 in loop1, is a crucial site for channel gating and inhibitor (carbenoxolone, CBX) interactions that block PANX1^23,24^. Incidentally the PANX3 has a branched hydrophobic aminoacid, isoleucine (I74), at this position. In this study, we present the electron cryomicroscopy (cryo-EM) structures of human Pannexin3 (PANX3) and mutants of human Pannexin1 (PANX1) to observe effects on channel organization, pore radii and effects on ATP interactions. The PANX3 structure displays a constriction point below the pore that facilitates the formation of a second vestibular cavity in PANX3. We observe that the congenital mutant R217H (PANX1_R217H_) in TM3, in comparison with PANX1_WT,_ results in the allosteric modulation of the PANX1 pore to constrict it further and significantly alter its conductance properties. Cationic residue substitutions at distinct locations within the channel vestibule ranging from the N-terminus to the pore lining residues including K24, R128, R217H and W74R/R75D into PANX1 yields altered ATP analog interactions. We study structure of PANX3 in comparison to other isoforms and demonstrate alterations in channel gating in PANX1 as a consequence of critical mutations in PANX1.

## Results

### Pannexin3 has distinct structural features in comparison to other isofoms

The PANX3 shares 42% sequence identity with PANX1(Figure S1) and 24% identity with PANX2. To understand the disparity among PANX isoforms, we purified the full-length human PANX3 and eludicated the structure to a resolution of 3.9 Å by cryo-EM in the presence of ATP (1mM) and K^+^ (100 mM) (Figure1A, B, Figure S2, Table S1). Similar to PANX1, high extracellular potassium is also speculated to open the PANX3 channels^29^. We used 100 mM K^+^ during final purification of PANX3 in an attempt to capture an open conformation of the channel. The density at the pore and transmembrane helices was sufficient to model a bulk of the side chains in these regions (Figure S3). The intracellular helices (160-185) and the C-terminus (373-392) lack clear densities, likely due to inherent flexibility in this region. Density for bound ATP molecules that would allow unambiguous assignment, was not observed. The PANX3 retains the heptameric oligomer assembly that can be partitioned into an extracellular domain (ECD), transmembrane domain(TMD), and intracellular domain(ICD), similar to PANX1_WT_ and PANX2 (Figure 1A, B). Unlike other large-pore ion-channels like CALHMs^30^, the PANX isoforms do not display heterogenous oligomeric associations^31^. The PANX3 channel is 8 Å wider than the PANX1 at the cytosolic face with similar transmembrane length in the ordered regions of the channel (Figure 1C, lower panel). The channel width of PANX3 at the extra and intracellular faces is similar to PANX2 (Figure 1C).

**Figure. 1.**
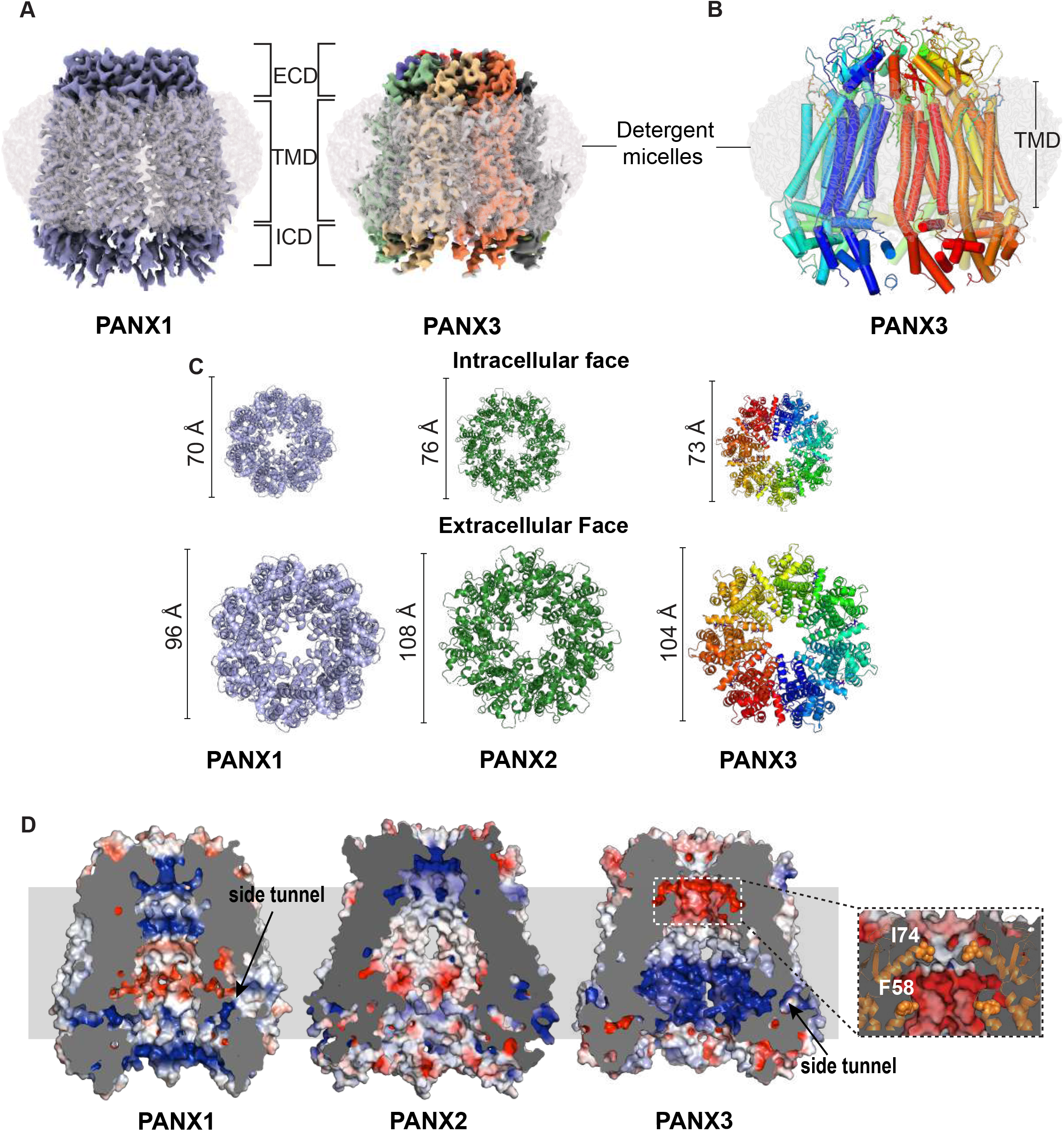
Structural comparison of PANX1 and PANX3. **(A)**, The cryo-EM map for PANX1(blue) and PANX3(monomers colored individually) viewed parallel to the membrane plane and surrounded by the detergent micelle (gray). **(B)**, The modeled structure of PANX3 diplaying heptameric organization and embedded in a detergent micelle that marks the transmembrane boundaries. **(C)**, Top and bottom views of PANX1, 2 and PANX3 exhibiting differences in the width in PANX isoforms at the extracellular and intracellular faces of the channel. **(D)**, Sagittal section of surface electrostatics of PANX1 (PDB ID: 6WBF), PANX2 (PDB ID:. 7XLB) and PANX3 colored according to potential from -5(red) to +5(blue) (*k*_*B*_*Te*_*c*_^*−1*^) viewed parallel to the membrane plane. The inset shows the position of the residues I74 and F58 (orange spheres) forming first and second constrictions, repectively in PANX3.

The topology of PANX3 protomers is similar to PANX1_WT_ with differences in the TM1 and the extracellular loops. The topology comprises four transmembrane helices and two extracellular loops (EL). Two disulfide bonds (SS1 and SS2) stabilize the extracellular domain, C66-C261(SS1) and C84-C242 (SS2), that form between EL1 and EL2 (Figure S4A, B). The superposition of two protomers (PANX1 vs. PANX3) results in a root mean square deviation (rmsd) of 3.2 Å for the 302 Cα atoms. A two stranded β-sheet is formed in PANX3 between EL1 and TM2 which has greater similarity with PANX2 structure in comparison to PANX1 that lacks this motif. There is also a significant alteration of surface charge within PANX3 compared to PANX1_WT_ and PANX2 near the surface lining the aqueous channel vestibule (Figure 1D).

N-linked glycosylation at the N255 position in the EL2 of PANX1 was implicated in preventing the formation of gap junctions^24,32^. A substitution at this site (N255A) led to the formation of a mixture of gap junctions and hemichannels^24^. In the PANX3 structure, we observed density for N-acetylglucosamine (NAG) at the predicted glycosylation site (N71) in the first extracellular loop that is much closer to the pore, in comparison to PANX1 (Figure S4C-D). The N-glycosylation in PANX2 is also localized to the EL1 region at N86 position that coincides with PANX3 N-glycosylation site (Figure S1) although the structure of PANX2 does not reveal the site due to a likely disorder in the vicinity^27^.

In PANX3 we did not observe density for residues 1-24 in N-terminus. It is therefore unclear if the N-terminus lines the pore and plays a role in maintaining the rigidity of the transmembrane domain of the heptamer similar to PANX1. A comparison with the Alphafold2 model of PANX3 with the experimental model in this study indicates that the N-terminus faces the cytosol (Figure S5A). It constricts the channel from the cytoplasmic side, unlike PANX1, where the N-terminus lines the pore and interacts partially with the adjacent protomer^24^. This disparity among isoforms has been observed in CALHM1 and 2, where the N-terminus of CALHM2 is directed towards the cytosolic side of the channel^33,34^. Towards the C-terminus, we did not observe the density for the last twenty residues (373-392). The C-terminus is comparatively shorter than PANX1 and lacks a caspase cleavage site to facilitate ATP release, as observed in PANX1. However, ATP release has been observed in PANX3 in presence of high extracellular potassium^29^. Consequently, the short C-terminus in PANX3 may not play a role in channel opening, and PANX3 may have alternate mechanisms for channel-gating. However, further experimental investigation is required for understanding the role of C-terminus in PANX3.

### PANX3 vestibule is organized into two compartments

The residue lining the constriction point in PANX1, W74, is replaced by isoleucine or valine in different orthologues of PANX3 (Figure 2A, B, Figure S5C). The PANX3’s pore is lined by two residues, I74 and R75, with a width of 13.2 Å at the narrowest point created by I74 (Figure 2B, C). The primary constriction created by the W74 residue in PANX1_WT_ results in a pore diameter of 12 Å. The presence of I74 in PANX3 instead of W74 widens the pore to a radius of 13.2 Å. The cation-π interaction between W74 and R75 in PANX1_WT_ is eliminated in PANX3 due to the replacement of W74 with I74. R75 of one protomer interacts with D81 of the other protomer through a salt bridge. Residues in the same position in the pore in PANX2 are substituted with R89 and D90 that are drastically different from residues observed in PANX1 and PANX3. R89 in PANX2 is suggested to form highly cationic pore at the channel entrance (Figure 2D-H, Figure S5B)^27^.

**Figure. 2.**
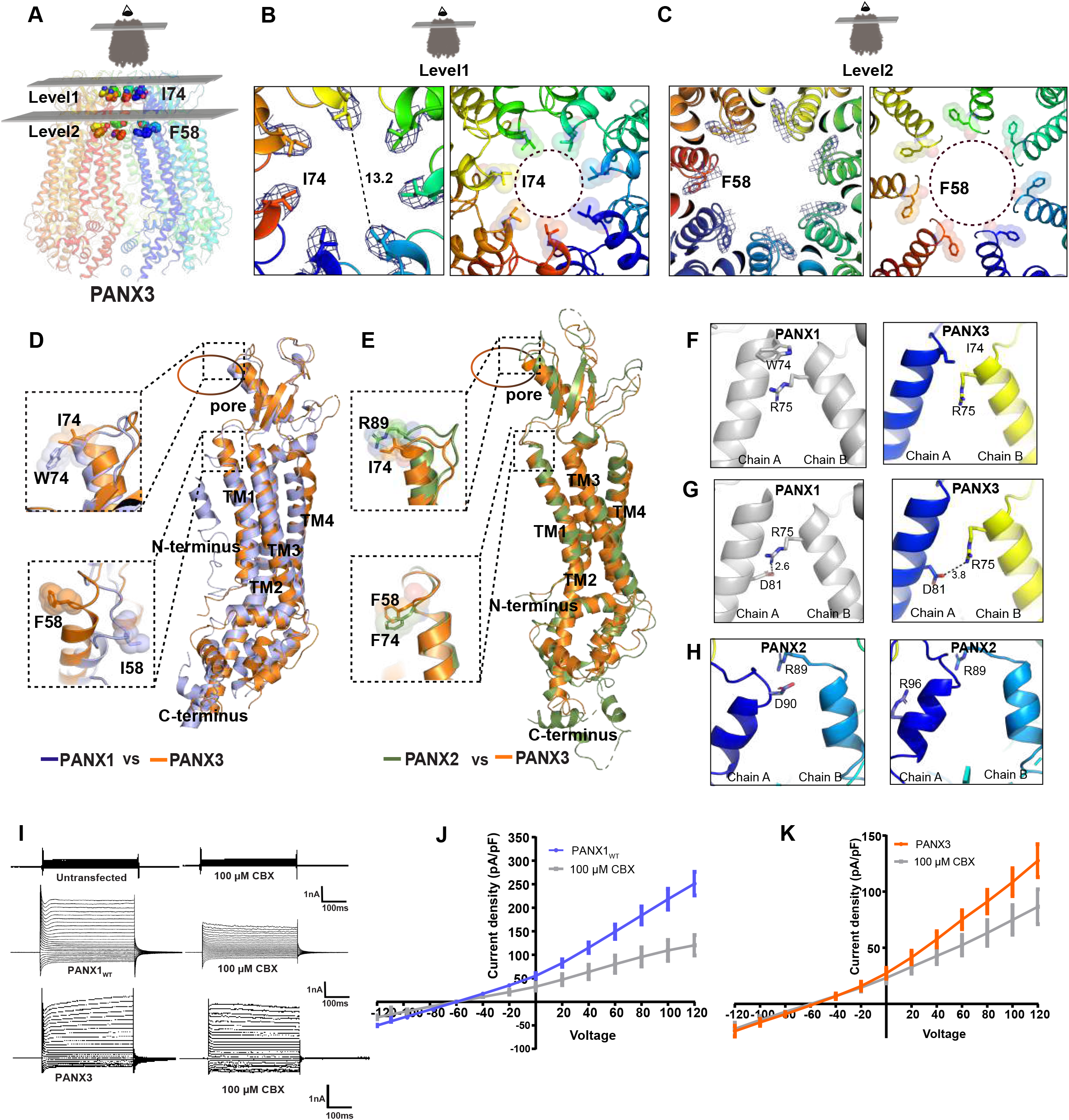
PANX3 vestibule is organized into two compartments. **(A)**, Transverse section in PANX3 at two levels displaying the positions of the two constrictions. The first constriction is formed by I74 in PANX3, and F58 in PANX3. **(B)**, A close-up of the first constriction seen from above in PANX3 (right panel) formed by I74. The density for the residue (I74) at 7.5 σ is shown(left panel). **(C)**, A close up of the second constriction formed by F58 in PANX3. The density for the residue (F58) at 7.5 σ is shown. **(D)**, Superposition of PANX1 and PANX3 displaying the position of residues 58 and 74 with a rmsd of 3.2 Å for 302 Cα atoms, **(E)**, Superposition of PANX2 (PDB ID: 7XLB and PANX3 exhibits the differences in the pore residues, R89 and I74 in PANX2 and PANX3 respectively with a rmsd of 5.4 Å for 288 atoms aligned; F74 in PANX2 acquires similar position as F58 in PANX3 **(F)**, Cation-pi interaction between W74 and R75 in PANX1 (PDB id 6WBF, left panel) is lost in PANX3 (right panel) **(G)**, Hydrogen bond interaction between R75 and D81 in PANX1 (left panel) is also observed in PANX3 (right panel). **(H)**, Pore residue(R89, D90) in PANX2 do not form any interactions with neighbouring residues unlike PANX1 and PANX3 **(I)**, Representative traces for whole-cell current for HEK293 untransfected cells(mock) and HEK293 cells expressing PANX1_WT_ and PANX3 with and without CBX (100μM) application. **(J-K)**, Current density-voltage plot for PANX1 and PANX3 in presence and absence of CBX. Each point represents the mean of n=4-5 individual recordings, and the error bar represents SEM.

Similarly, S70 and Q76 between two protomers of PANX3 form a hydrogen bond resulting in interprotomeric interactions leading to a stable heptamer. Apart from these interactions, there seem to be minimal intersubunit interactions across the transmembrane region between the protomers. The gap between the TMs 2 and 4 of adjacent protomers just beneath the ECD is occupied by lipid-like density into which we have modeled a phospholipid, 1-palmitoyl-2-oleoylphosphatidylethanolamine (POPE), for each protomer (Figure S4E). We could also observe a gap between subunits that is large enough to allow the passage of ions. The presence of a side-tunnel suggested by Ruan et al. in PANX1_WT_ is more prominent in PANX3 with a 6.9 Å separation between TM2 and CTH1 compared to 6.3 Å in PANX1_WT_, suggesting that the lateral portal hypothesis may also hold true for PANX3 (Figure S5D). The region is occluded with a distance of 2.6 Å between TM2 and CTH1 in the case of PANX2 and is unlikely to support the entrance of ions from this portal in PANX2 (Figure S5D).

The structural comparison of PANX3 with PANX1_WT_ reveals an additional constriction in PANX3 below the primary channel pore. The residues at the end of TM1, 58-60 comprising residues F58, S59, and S60 forms a prominent second constriction at the neck region in PANX3 compared to PANX1_WT_ (Figure 2A-D). The linker between TM1 and TM2 adopts a clear α-helical conformation in PANX3 and PANX2, instead of a loop observed in PANX1_WT_, constricting the vestibule in the region, thus allowing PANX3 to have an additional vestibule beneath the pore (Figure 2A-D). The residues F58-S59-S60 line the second constriction point facing the pore in PANX3 and are part of the PANX3 sequence that is variable between PANX1, 2 and 3 (Figure S5E). In contrast, I58 residue in PANX1_WT_ participates in hydrophobic interactions between TMs 1 and 2. Despite sequence variation between PANX2 and PANX3, the F74 in PANX2 forms a similar motif that can allow the demarcation of the vestibule into two regions even in PANX2 isoform (Figure 2E). The diameter at this constriction point is 21 Å compared to 30 Å in PANX1_WT_ and demarcates the boundary between the anionic surface of the upper compartment compared to the amphiphilic lower compartment in PANX3 (Figure 1C, D). An annulus of seven uncharacterized densities is observed at this constriction point that might further contribute to restricting access at this region(Figure S5F). We performed a binding analysis of ATP-γS with PANX3 using microscale thermophoresis (MST). Both ATP and ATP-γS displayed similar affinities when tested using PANX1_WT_ that encouraged us to use the non-hydrolysable analogue of ATP for MST binding assays (Figure S6A, B) The affinity of ATP analog ATP-γS for PANX3 was determined to be 75 μM compared to the 13 μM affinity observed with PANX1_WT_ (Figure S6C). The ATP-γS binding studies suggest a weaker affinity of PANX3 for ATP-γS compared to PANX1. Although we observe a lower affinity of ATP-γS for PANX3, it is difficult to speculate the factors affecting the affinity, as a clear ATP binding site is yet to be detected in large pore ATP channels such as CALHM, connexins and pannexins. Moreover, inconsistencies in our ATP release experiment make it difficult to understand the correlation between binding affinity with ATP release activity of the channel (data not shown).

The different pore-lining residues observed in PANX1 and PANX3 may dictate isoform-specific differences in channel properties as observed in connexins^35^. Despite earlier unsuccessful attempts to record currents from PANX3, we could measure currents in PANX3 expressed in HEK293 cells^22,23^. Current density measurements and I-V curves for PANX3 suggest that it has a similar voltage sensitivity as PANX1 and elicits outwardly rectifying currents at positive potentials (Figure 2I-K). We observe weaker CBX sensitivity in PANX3, in comparison to PANX1(Figure S7A). The ECD region, particularly loop1, surrounding the pore was previously implicated as being important for CBX interactions^23^. The substitutions among pore lining residues observed in PANX3 could be a reason for the reduced sensitvity to CBX. The ability of CBX to interact with PANX3, albeit weakly, despite a wider pore, reinforces the idea that CBX interactions at extracellular domain can modulate channel activity instead of directly acting as an asymmetric pore blocker. Alternate uncharacterized sites for CBX interactions can exist within PANX isoforms given its sterol like chemical structure that modulates channel activity. This is reinforced by the lack of saturation in CBX interactions when tested using binding experiments (Figure S6D). The structural and the vestibule surface electrostatic differences observed between the three isoforms PANX1, 2 and 3 can therefore influence their respective channel activities and, consequently, their physiological properties. We therefore evaluated the effects of multiple substitutions of cationic residues including a PANX1 congenital mutant on channel properties along the vestibule.

### Pannexin1 congenital mutant alters channel properties

We elucidated the structure of PANX1 R217H substitution to characterize the effects of this congenital mutant and to understand the basis of its defective channel properties. The arginine residue at the 217 position is highly conserved among vertebrate PANX1 orthologues (Figure S8A). The arginine was mutated to a histidine by site-directed mutagenesis. Consistent with the earlier reports^15^, we did not observe compromised expression of the PANX1_R217H_ channel compared to PANX1_WT_ (Figure S8B). The protein was purified to homogeneity, and the structure was determined by cryo-EM to a resolution of 3.9 Å (Figure S9B, Table S1). The model was built using the previously reported structure of PANX1_WT_ (PDB ID: 6WBF). The superposition of PANX1_WT_ and PANX1_R217H_ mutant yielded a rmsd of 1.2 Å for 288 Cα atoms.

We compared the structure of the mutant channel with the PANX1_WT_ to analyze the local and global variations. The structure resembles PANX1_WT_ globally, with changes in extracellular domain (ECD), extracellular helix 1 (EH1), and the intracellular domain (ICD). Extracellular loops and EH1 have an average shift of 1.5-2 Å away from the pore, which along with the slight TM1 movement, is most likely responsible for inducing the change in the W74 position by moving the tryptophan indole ring towards the pore, thereby shortening the pore radius by 4 Å (Figure 3A, B). The R75 residue interacts with the D81 of the adjacent subunit through a salt bridge, consistent with PANX1_WT_.

**Figure. 3.**
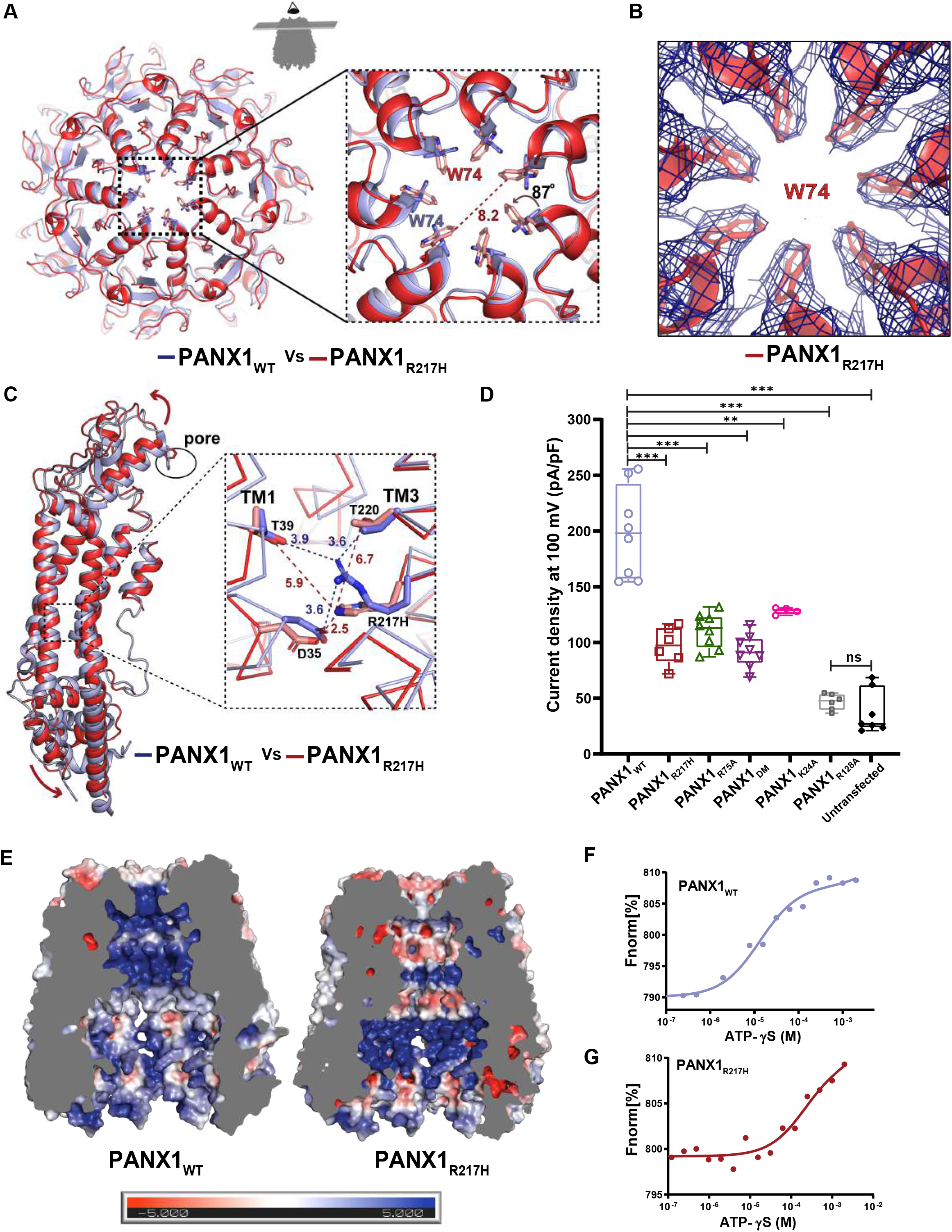
PANX1 germline mutant, PANX1_R217H_ alters channel properties. **(A)**, A cross-section of superposed structures of PANX1_WT_ (blue) and PANX1_R217H_ (red); the inset shows the altered conformation of the residue (W74) at the extracellular entrance of the pore. The dashed line represents the reduced pore cross-section distance in PANX1_R217H_(red); the W74 rotates by 87° at the χ^2^ torsion angle, reducing the pore radius by 3.8 Å compared to PANX1_WT_ (Supplementary Movie1). **(B)**, The density for W74 in the mutant PANX1_R217H_ channel. **(C)**, The structural superposition of PANX1_WT_ and PANX1_R217H_ exhibits a disrupted hydrogen-bond network, owing to the mutation, displayed in the inset. For clarity, only one subunit is shown, and arrows indicate the direction of the movement of the mutant in comparison to PANX1_WT_. **(D)**, Current density is plotted for the PANX1_WT_(n=8) and the mutants, PANX1_R75A_(n=8), PANX1_R217H_(n=6), PANX1_DM_(n=8), PANX1_R24A_(n=4) PANX1_R128A_(n=6) and untransfected(n=6) the error bar represents SEM. n represents the number of cells used for independent recordings; a two-tailed unpaired t-test is used for calculating the significance, ***p < 0.001; n.s., not significant. **(E)**, Surface representation of PANX1_WT_ and PANX1_R217H_ according to the electrostatic surface potential from -5(red) to +5(blue) (*k*_*B*_*Te*_*c*_^*−1*^) viewed parallel to the membrane plane. The residues in the full length PANX1 structure (PDB ID:6WBF) were deleted to match the residues in the modeled PANX1_R217H_ structure, to compare the electrostatic surface potential between the structures owing to the similar residues., **(F)**, Binding affinity for PANX1_WT_ with ATP-γs was estimated as 13±3 μM. **(G)**, Weak apparent binding affinity of the PANX1_R217H_ with ATP-γs was determined to be 210±141 μM compared to 13±3 μM of PANX1_WT_. The experiments are done in triplicates (Figure S14)

As the mutation of R217H is in TM3, we investigated the residues in the vicinity of R217 within a 4 Å radius. The D35 in TM1 interacts with R217 (3.6 Å) in PANX1_WT_ and forms a H-bond. R217 also interacts with T220 in TM3 and T39 in TM1 through hydrogen bonds (Figure 3C). Mutating arginine to histidine disrupts the H-bond interaction network within TMs 1 and 3. The H-bond interaction of T220 in TM3 and T39 in TM1 with R217H is disrupted due to the shorter side chain of histidine compared to arginine. The G44 in TM1 acts as a hinge point that facilitates the TM1 bending and displacement as a consequence of R217H substitution. The displacement observed in TM1 translates to an outward movement of ECD that allows the W74 to flip towards the pore (Figure 3A, B).

In comparison to the PANX1_WT_, the W74 shifts by a χ2 torsion angle of nearly 80° towards the pore leading to substantial closure of the pore diameter in PANX1 channels and affecting channel properties (Figure 3A, Movie S1). A comparison of the current density(pA/pF) of R217H to PANX1_WT_ reveals weakened channel activity (Figure 3D). The average current density for PANX1_WT_ was twofold higher than PANX1_R217H_, at a positive voltage of 100 mV suggesting partial closure of the pore (Figure 3D, Figure S7B). Although there was a significant decrease in the current density in the R217H mutant, the mutation did not have significant effect on the CBX binding, as we could observe the inhibition of PANX1_R217H_ currents by CBX similar to PANX1_WT_ (Figure S7A, 10). A comparison of the conductance density (nS/pF) changes to voltage reveal highly weakened channel conductance in response to increasing voltage in comparison to PANX1_WT_. A comparison of normalised conductance to voltage for PANX1_R217H_ and PANX1_WT_ exhibits altered voltage sensitivity (V_50_)of the mutant channel from around 40 mV for the PANX1_WT_ to over 100 mV indicating a loss of channel conductance as a consequence of R217H mutation (Figure S7C, D, E).

Binding studies using MST reveal a decreased ATP-γS binding in the PANX1_R217H_ mutant compared to PANX1_WT_ (Figure 3F, G). As observed in PANX1_R217H_, altering charged residues in the vestibule affects the electrostatic surface of the PANX1 vestibule significantly. Although the R217H substitution is in the TM3, we detect structural shifts in the ECD indicating allosteric effects of this congenital mutant. Such allosteric effects are observed in the case of disease-causing mutants where the mutation site is far from the observed structural changes and affects their functional properties^36^. It was proposed in an earlier study that the PANX1_R217H_ interactions with the C-terminus can cause the altered channel properties in this congenital mutant^37^. Given the allosteric effects of R217H on the pore diameter observed in this study, long-range effects in C-terminus could also drive some of the properties observed with PANX1_R217H_. However, given the absence of a structured C-terminus in PANX1 structures determined thus far, the influence of the C-terminus on PANX1_R217H_ structure would be speculative at this juncture.

### Substitutions in PANX1 vestibule alter ATP interactions and channel behavior

Altering positively charged residues in PANX1 can alter the properties of the channel as observed in PANX1_R217H_^24-26^. In the context of ATP binding, we explored the effects of substituting other conserved positively charged residues along the vestibule, like K24, R29 (N-terminus), K36 (TM1), R75 (EH1, pore) and R128 (TM2), that face the vestibule of the PANX1 channel and can influence ATP interactions (Figure S6E). The substitution of R29A and K36A resulted in excessive cell death and purification of these PANX1 mutants could not be done. The ATP-γS interaction with K24A substitution has a six fold loss of ATP-γS affinity (*K*_d_ of 75 μM) compared to PANX1_WT_. On the other hand, R128A (TM2) substitution reveals a complete loss of ATP-γS binding, indicating the importance of this region for PANX1 function (Figure S6F, I). Moreover, R75 was suggested as a putative ATP binding site through alanine scanning mutagenesis^38^. We, therefore, mutated R75 to alanine and checked the binding affinity of PANX1_R75A_ with ATP-γS. The MST data suggests a decrease in the binding affinity with PANX1_WT_ (Figure S6G). Although we observe that R75 contributes to ATP-γS binding, R75 does not act as the sole residue for ATP binding as mutations in other positive residues also led to decreased ATP-γS binding (Figure S6F-K). In all likelihood, multiple residues are involved in ATP interactions, and mutating these residues individually reduces ATP binding but does not abolish it entirely.

To further investigate the effects of these mutations, we performed whole cell patch clamp studies. The R128A PANX1 behaved similar to the untransfected cells and the loss of ATP-γS binding in R128A could be a result of improper oligomerization, as observed through the widened size-exclusion chromatography profiles (Figure S11G). The N-terminus mutant K24A, exhibited weaker current density compared to the wild type in electrophysiology studies (Figure 3D, FigureS7B). Moreover, electrophysiology data shows a two-fold decrease in the current density in PANX1_R75A_ compared to PANX1_WT_, although the whole cell expression for both mutants is comparable to the PANX1_WT_, suggesting that the substitutions in the vestibule and the pore can affect the function of the PANX1 channel (Figure S7B, S8C).

However, normalized G-V curve for R75A is comparable to wild type and reveals that the substitution of a pore residue, R75, does not affect the voltage sensitivity of the channel (Figure S7D-E) consistent with the findings with ZfPANX1^39^. Similar consequences to arginine substitutions in the pore are observed with LRRC8A, where the channel maintains conductance with arginine substitution but loses anion selectivity^40^.

Single substitutions of W74 and R75 in the pore region also cause an incorrect assembly of PANX1^24^. In order to observe if dual substitution of both the residues (W74, R75) by charged amino acids could compensate for this behavior, we created a PANX1 double mutant (PANX1_DM_) by substituting W74 and R75 with arginine and aspartate, respectively. These substitutions are similar to the pore residues of PANX2 revealed by multiple sequence alignment (Figure S1). Surprisingly, we obtained a minor fraction of well assembled PANX1_DM_ channel whose structure of PANX1_DM_ was elucidated to a resolution of 4.3 Å (Table S1, Figure S9C, 11C). The model was built using the previously determined structure of PANX1_WT._ Although the overall resolution is low, the densities for the pore residues were sufficient for positioning aminoacid sidechains at this location (Figure S8D). Whilst the global structure resembles the PANX1_WT_, the pore diameter was reduced to 7.9 Å due to the presence of R74 side chain facing the pore that renders it highly cationic as compared to PANX1_WT_ and the pore diameter is comparable to PANX2 (8.8 Å) (Figure S8C-F)^27^. Using electrophysiology, we explored the effects of these structural changes on channel physiology (Figure S7). The data revealed a two-fold reduction in the current density in PANX1_DM_ than PANX1_WT_, which can be correlated with the reduction in pore radius (Figure S7B, S8D). The ATP-γS binding data shows a weakened *K*_d_ value of 78 μM in PANX1_DM_ compared to ATP-γS binding affinity for PANX1_WT_ (13 μM), suggesting that ATP interactions are partly affected by these pore substitutions (Figure S6H). We observe a decreased inhibition of PANX1_DM_ currents with CBX compared to PANX1_WT_ since the primary binding site for CBX, W74, was substituted to an arginine (Figure S7A). It is rather interesting that CBX retains minimal interactions with the PANX1_DM_ despite major substitutions in the pore. This further indicates that presence of alternate sites of interaction could modulate channel gating. The conductance density shows a fall in channel conductivity for PANX1_DM_ although normalised G-V curve plotted for the double mutant and the pore mutant R75A displays an unaltered voltage sensitivity (Figure S7D,E). Thus, we can infer that the pore substitutions do not influence the voltage-dependent conductance property of the channel.

In conclusion, our structural findings suggest that pore radii are inherently linked to residues lining this constriction and can have distant structural effects. It is observed in this study that a substitution, R217H, that alters the H-bond network within TM regions has dramatic effects on the pore radius leading to some of the observed physiological consequences.

## Discussion

In this study, we report the previously unknown structure of PANX3, an isoform of PANX1. PANX3 shows heptameric oligomer assembly consistent with PANX1 but differs in tissue localization^17^. We observe a voltage dependency similar to PANX1 as it responds to positive voltage but has a weaker affinity for ATP-γS. The pore size of nearly 13.2 Å indicates the conformation of PANX3 is likely to be an open state that can allow the passage of multiple ions, dyes, and possibly ATP.

A unique feature of PANX3 is the second vestibule in the neck region, which has not been previously observed in any other large-pore channel. The region F58-S59-S60 at the neck faces the pore, forming a ring and demarcating PANX3 vestibule into two compartments (Figure. 2A). Although the pore is quite wide around F58 in PANX3 structure, the flexibility in the side chain could dynamically regulate the size of this constriction point. This could aid in the regulation of PANX3 channel activity, particularly in the context of PANX3 lacking a long C-terminus that is caspase-sensitive, unlike PANX1. The presence of hydrophobic residues at the neck has been observed in ion channels like Bestrophin^41^ and pentameric ligand-gated ion channels (pLGICs)^42^. Hydrophobic constrictions in ion channels can aid in the creation of barriers to ion flow and alter the wettability of the pore^43^.

The cryo-EM structure of congenital mutant, PANX1_R217H_, reveals a narrower pore in comparison with PANX1_WT_ as a consequence of the rotation of W74 at its χ2-torsion angle by ∼80° and partially closes the pore (Movie S1). This is facilitated by the outward movement of the extracellular domain away from the pore as a consequence of the R217H substitution in TM3 that locally destabilizes the H-bond network in the TM region. The weakened conductance behavior of this mutant signifies its inability to respond to a voltage similar to PANX1_WT_. The transition in pore diameter observed in this study as a consequence of the TM substitution indicates the presence of allostery within PANX1 channels that also correlates with the reduced ATP efflux observed with this congenital mutant of PANX1^15^. The substitution of charged residues like R217H, R128A, and K24A around the vestibule also alter the surface electrostatics of the internal vestibule of PANX1, which can affect the channel function.

The double substitution of residues in PANX1 pore, W74R/R75D (PANX1_DM_), also exhibits a smaller pore than PANX1_WT_ consistent with PANX2. The PANX1_DM_ mimic resembles LRRC8A, where the constriction is lined by a ring of positively charged arginine residues and plays an essential role in anion permeability^40^. In other isoforms of LRRC8, the arginine is replaced by leucine and phenylalanine; a similar hydrophobic substitution can also be seen in PANX3. The closure of the pore is correlated to the weakened current density and a reduced ability of CBX to interact with the pore of PANX1.

An analysis of the pore radii of all the constructs, generated through the program ‘Hole’, reveal differences in the channel pore of the mutants compared to PANX1 _WT_^44^. The main distinction lies in the width of the pore entrance depicting mutants’ smaller size than PANX1_WT_. The smallest Van der Waal radii found were 2.4, 5.3, and 4.2 Å for PANX1_R217H_, PANX3, and PANX1_WT,_ respectively, at the pore entrance (Figure 4A-D).

**Figure. 4.**
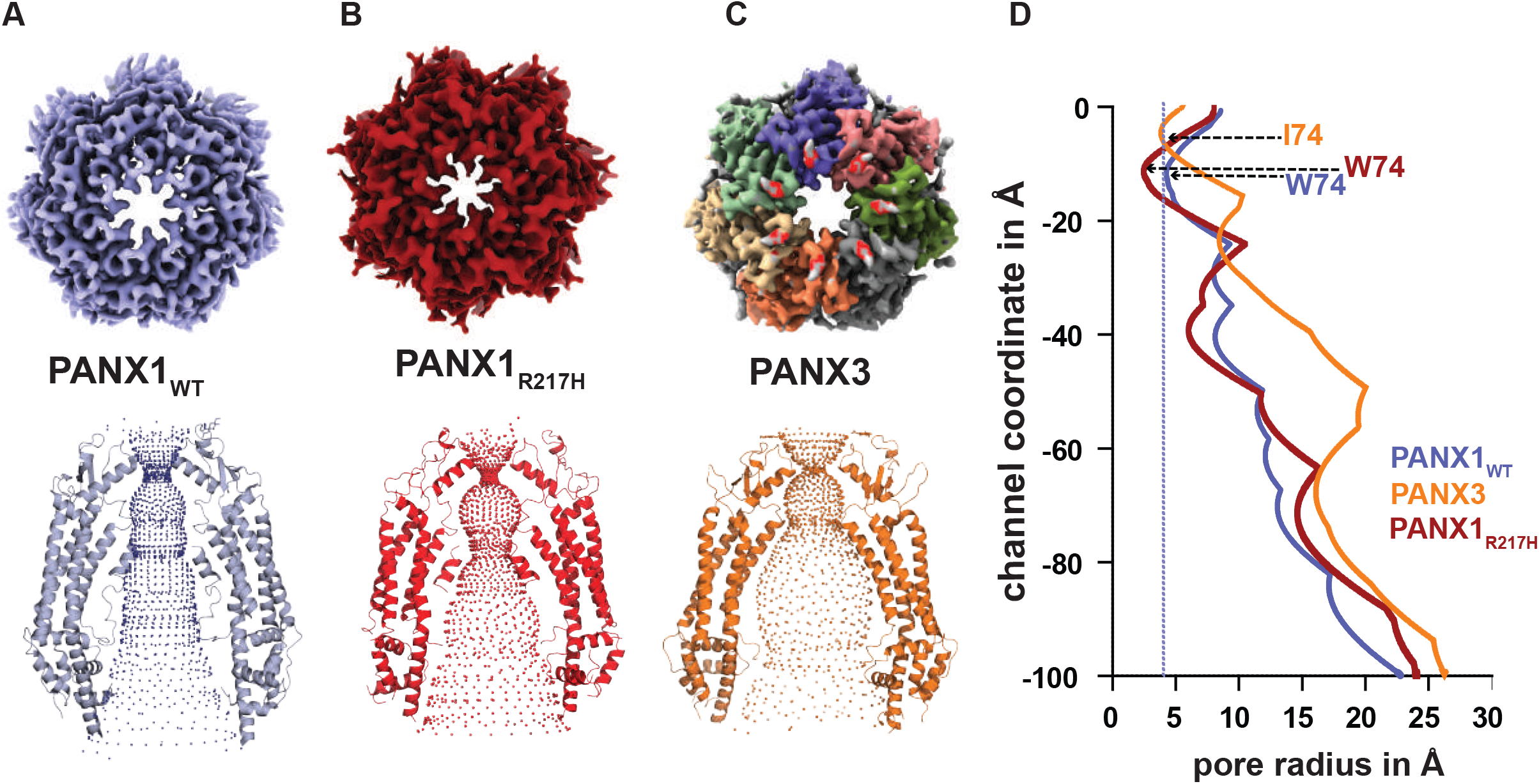
Pore profiles generated by hole program. The minimum radius is observed at the first constriction in PANX isoforms. **(A)**, The minimum radius is 5.3 Å for PANX3 at the constriction formed by I74. **(B)**, The PANX1_WT_ has a minimum radius of 4.2 Å formed by W74. **(C)**, The constriction points are created by W74 in PANX1_R217H_. The constriction point is 2.4 Å in PANX1_R217H_. (**D)**, 2D representation of the hole profile. The line represents the minimum radius for PANX1_WT_ at 4.2 Å formed by the first constriction (W74) in PANX1 channels.

Previous structures of PANX1_WT_ were determined by multiple groups in different conditions. For instance, high K^+^ has been used extensively to open the channel^24,25^, and C-terminus cleavage has also been shown to increase the current density in whole-cell recordings^24^. However, all the previously determined structures have identical channel organization and similar positions of pore-lining residues^24,25^ (Figure S12B-D). In this study, as a consequence of a congenital mutant and pore substitutions in PANX1, we could observe changes to the pore radius that are consistent with the altered channel behavior of PANX1_R217H_ and also highlight the pore constriction induced by a double substitution of W74R/R75D.

In conclusion, the study gives insights into the novel aspects of PANX3 architecture and demonstrates the comparison of pore radii with PANX1 and PANX2 (Figure 5). The presence of long-range effects of a congenital mutant further underscore the interesting structural and functional diversity among pannexins.

**Figure. 5.**
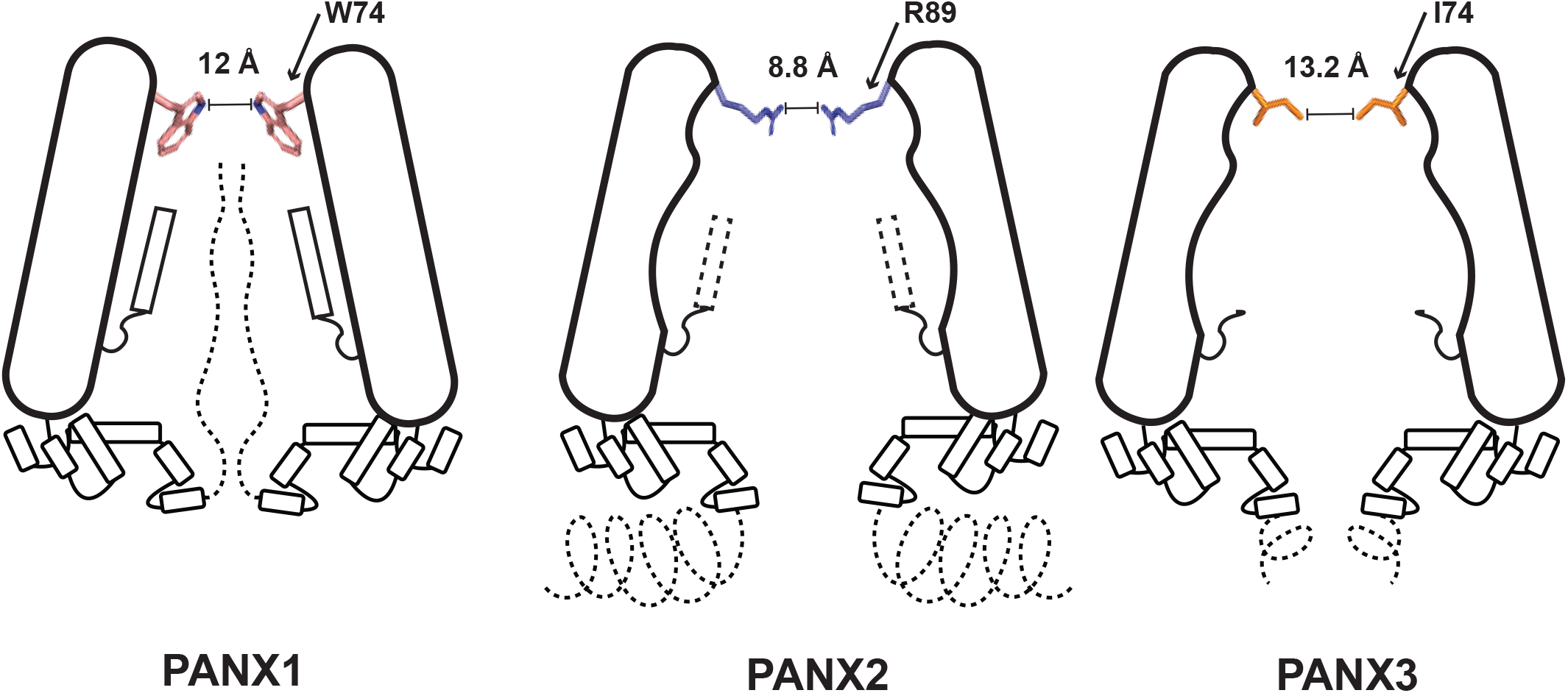
Schematic of PANX isoforms comparison. The schematic of the three PANX1 isoforms displays the differences in their structural organization at the pore due to altered residues within the pore at the extracellular domain. The distance between residues at the constriction point are provided.

## Acknowledgments

Research in this manuscript is funded by the MHRD-STARs program individual grant to AP(MoE/STARS1/167). The work in the project was partly funded by the WellcomeTrust/DBT-India Alliance intermediate (IA/I/15/2/502063) and senior (IA/S/22/1/506242) fellowships to AP. NH is a PhD student funded by the DBT-JRF program (DBT/2017/IISc/877) at IISc. AA is a PhD student funded by the IISc-GATE fellowship. SM was a graduate student at IISc funded by the DST-INSPIRE fellowship. SP was a postdoctoral fellow of the DBT-RA program in biological sciences. APB is an early-career fellow of the DBT-Wellcome Trust India Alliance. We thank Sucharita Bose and the National Electron Cryo-Microscopy facility, BLiSc, Bangalore (DBT/PR12422/MED/31/287/2014), for the screening of grids and data collection. We would like to acknowledge the advanced cryo-EM facility, IISc funded through the DBT-BUILDER program (BT/INF/22/SP22844/2017) and DST-FIST program (SR/FST/LSII-039/2015) for preliminary data collection. Computational support from the high-performance computing facility “Beagle” setup from grants by a partnership between the DBT, India, and the Indian Institute of Science (IISc-DBT partnership program) is acknowledged. The authors would like to acknowledge the DBT-IISc partnership program phase II and the DST-FIST program support for research.

## Author Contributions

NH performed the molecular cloning, ATP binding experiments, optimized purification, sample preparation, grid freezing, performed data processing, refinement of PANX structures and data analyses. AA performed and analysed all the electrophysiology experiments under guidance from SKS. SP initiated the cloning of PANX1 gene and optimized heterologous expression. SM performed initial electrophysiology measurements. APB aided in the preliminary experiments with PANX3. VKR performed the data collection, initial data processing and optimization of grid freezing for PANX1. AP designed and planned the project and provided inputs for structure refinement and data analyses. NH and AP wrote the manuscript with inputs from all authors.

## Methods

### Plasmids and cloning

Full-length human PANX1 (UniprotID-Q96RD7) and PANX3 (UniprotID-Q96QZ0) were synthesized by Geneart (Invitrogen). The synthesized genes were subcloned into a pEG-BacMam vector between EcoR1 and Not1 restriction sites with a C-terminal TEV protease site (ENLYFQ) followed by enhanced green fluorescent protein(eGFP) and 8x His-tag^45^.

The site-directed mutagenesis was used to generate the mutants by the mega primer-based whole plasmid amplification method. The primers were designed using SnapGene and synthesized by Sigma. The mutations were verified by Sanger sequencing.

### Transfections and Florescence-detection Size Exclusion Chromatography

HEK293S GnTI^−^ cells were seeded at a density of 1×10^6^ cells in DMEM medium with 10% FBS. The cells were transfected using lipofectamine 3000 (Invitrogen) according to the manufacturer’s protocol. For fluorescence-detection size exclusion chromatography (FSEC), cells were harvested after 36 hours and were solubilized using 200 μl of 25 mM Tris pH 8.0, 100 mM KCl, 1% glycerol, and 10 mM glycodiosgenin (GDN). The solubilized cells were spun down at 66,000g for 1 hour, and the supernatant was loaded onto the superose6 increase 10/300 GL column (GE Healthcare). GFP fluorescence (λ_Ex_=488 nm, λ_Em_=509 nm) was monitored to check the elution volume and the homogeneity of the protein (Figure S12A).

### Protein expression and purification

The protein was expressed using the *Bacmam* system for high-level expression^45^. In brief, *E*.*coli* DH10bac cells were transformed with a pEG-PANX1/3 vector. Positive colonies were selected based on blue-white screening, and bacmids were isolated and transfected into Sf9 adherent cells using cellfectin (Invitrogen) according to the manufacturer’s protocol. Four days post-transfection, cells were visualized for GFP fluorescence under a fluorescence microscope (the presence of green cells implies the presence of the virus). The virus was filtered through a 0.22 μm filter and harvested. Sf9 cells in suspension at a cell count of 2×10^6^ cells per ml were infected for the generation of the P2 virus. Four days after the infection, cells were spun down at 7000 g, and the supernatant was filtered and stored at 4°C, protected from the light.

HEK293S GnTI^−^ cells were grown at 37°C in Freestyle 293 medium(Invitrogen) supplemented with 1.5% FBS. Cells were infected with the P2 virus at a density of 2.5-3.0 × 10^6^ cells per ml. Twelve hours post-transfection, sodium butyrate was added at a final concentration of 5mM, and the temperature was reduced to 32°C. Cells were harvested 60 hours after infection for the enrichment of the membranes, and cells were sonicated for 15 minutes at 35% amplitude; the sonicated cells were centrifuged at 100,000×g for one hour. Membranes were flash-frozen and kept in -80°C till further use.

The expressed PANX1/3 was extracted from membranes using 10 mM glycodiosgenin(GDN), 25mM Tris pH(8.0), 100 mM KCl. The solubilized membranes were centrifuged at 100,000 × g for one hour. The supernatant was incubated with Nickel-NTA resin (Qiagen) equilibrated with binding buffer for two hours. The protein was eluted in 25 mM Tris pH (8.0), 100 mM KCl, 1% glycerol, 0.3 M imidazole and 100 μM GDN. The eluted protein was treated with TEV protease for C-terminal GFP and 8x-his-tag cleavage.

The GFP cleaved protein was further purified by size exclusion chromatography in 25 mM Tris pH(8.0), 100 mM KCl, 1% glycerol, and 50 μM GDN (SEC buffer) using Superose6 increase 10/300 GL column. The peak fraction was collected and used for grid freezing (Figure S11, 13).

### Cryo-EM sample preparation and data collection

The SEC purified PANX1/PANX3 was concentrated to 5 mg/ml using a 100 kDa cut-off concentrator (Millipore). The concentrated protein was centrifuged at 66,000 g for one hour before grid freezing. Quantifoil 300 mesh gold holey carbon grids (R1.2/1.3) were glow discharged in the air for one minute at 25 mA, and the protein (3 μl) was loaded on the grid in FEI vitrobot at 100% humidity and 16°C temperature. The grids were blotted for 3.5 s with a wait time was 10 seconds. For the PANX1_R217H_ mutant, samples were applied twice, and grids were blotted sequentially for 2.0 and 3.5 seconds, respectively. The grids were flash-frozen in liquid ethane and were stored in liquid nitrogen till further use.

For PANX3, ATP at a final concentration of 1 mM was added 30 min prior to the grid freezing. The ATP stock was prepared in SEC buffer (Tris pH 8.0).

Data sets were collected on a Titan Krios 300 keV (ThermoFisher) equipped with Falcon 3 or K2 direct electron detectors (Table S1). For PANX1_WT_ and PANX1_DM_, data was collected with the Falcon 3 detector, whereas for PANX1_R217H_ and PANX3, data was collected on the K2 detector in EFTEM mode with a Bioquantum energy filter and 20 eV slit width. The pixel size of 1.07 Å was used for both the detectors and the total electron dose for different constructs is mentioned in the table (Table S1).

### Cryo-EM data processing and model building

PANX1/3 structures were determined using the cryoSPARC version (3.20)^46^. The movies were imported and motion-corrected by patch motion correction, and the contrast transfer function (CTF) was estimated by patch CTF. After manually curating the images, the CTF-estimated micrographs were used for auto-picking by the blob picker. For manual curation, a cut-off of 6Å for CTF estimation and 1.06 for ice thickness was used to remove bad micrographs.

The auto-picked particles were extracted with a box size of 360 pixels (1.07 Å /pix) and were subjected to 2D classification; two rounds of 2D classifications were done to remove the junk particles. The selected 2D classes were subjected to ab-initio 3D classification. For the ab-initio modeling, the initial and final resolutions were kept at 12 Å and 8 Å, respectively, and the minibatch size of 1000 was used. The high-resolution class was used for non-uniform refinement^47^. The parameters for non-uniform refinement were adjusted to get a higher resolution map. The maximum alignment resolution was changed to the Nyquist limit, the number of extra passes was increased to 2, the initial dynamic mask resolution was changed to 14 Å, and the batch size was kept like that of ab-initio modeling. The map obtained from non-uniform refinement was subjected to local refinement; a separate mask for the local refinement was not generated, and a default mask provided by cryoSPARC was used. For PANX1/3 datasets, heterogeneous refinement did not produce any high-resolution map and thus was not used for the processing. PANX1 structure refinement was performed with C7 symmetry, whereas PANX3 was processed with C1 for the initial refinement; as C7 symmetry could be seen in the ab-initio 3D structure, C7 symmetry was applied during the non-uniform refinement to improve the resolution. PANX1 mutants were modeled using previously determined structures (PDB-ID: 6WBF). The desired residues were mutated, and the changes owing to the mutations were modeled in the cryo-EM density map in Coot^48^.

For PANX3, an Alphafold2 monomer was fitted in the map to make a heptamer in Chimera. The model and the map were aligned using autodock in Phenix. The model was built in the density in coot. All the structures were refined using PHENIX real-space refinement^49^. The workflow and the statistics for each construct are summarized in the extended figures and table (Figure S2, 9, Table S1).

### Binding studies with ATP-γs

Binding studies were done using microscale thermophoresis (MST). The protein concentration of 10 nM (calculated for the heptamer) was kept constant for all the studies.

The protein was labeled with 10 nM red Tris Pico dye, and Monolith standard treated capillaries were used to detect binding. A non-hydrolyzable ATP analog, ATP**-**γs, dissolved in SEC buffer, was used as a ligand for the binding studies. The ligand concentration was diluted (2x) for the study in 16-serial dilution, with 2 mM as the highest concentration. The final protein concentration was kept at 5 nM. All the experiments were done in triplicates. (Figure S14)

### Electrophysiology

HEK293 cells were maintained in DMEM F-12 Ham medium (Sigma) supplemented with 10% fetal bovine serum (GIBCO, heat-inactivated US origin) and 1 % antibiotic-antimycotic solution (Sigma) in a humidified incubator at 37°C with 5% CO_2_. The cells were passaged twice a week, and a fraction of the cells were plated onto 35 mm cell culture dishes (Thermofisher). PANX1 and its mutants and PANX3 were transiently transfected into HEK293 cells with an enhanced green fluorescent protein (eGFP) using the transfection agent Lipofectamine 2000 (Invitrogen). The cells expressing GFP were selected for patch clamp electrophysiological recordings after 24-36 hours of transfection.

PANX currents were activated and recorded in whole-cell mode using EPC 800 amplifier (HEKA Elektronik), Digidata 1440A digitizer (Molecular Devies), and pClamp 10 software. To elicit the currents, 400 ms voltage clamp steps were applied from a holding potential of -60 mV to test potentials of -120 to +120 mV in 10 mV increments. The currents were sampled at 20 kHz, and digitally low pass (Bessel) filtered at 3 kHz. The patch electrodes used for electrophysiology experiments were fabricated using borosilicate glass capillaries and had a resistance of 3-5 MΩ when filled with an internal solution. The pipette solution contained in mM) 150 Cesium gluconate, 2 MgCl_2_.6H_2_O, and 10 HEPES (pH adjusted to 7.4 with CsOH). The extracellular solution contained (in mM) 147 NaCl, 2 KCl, 1 MgCl_2_.6H_2_O, 2 CaCl_2_.2H_2_O, 10 HEPES, 13 Glucose (pH adjusted to 7.4 with NaOH)^15^. The effects of carbenoxolone (Sigma) were evaluated at a final bath concentration of 100 μM. The experiments were performed at room temperature (23°C). (Figure S10)

Electrophysiological data were analyzed using Clampfit, Microsoft Excel, and Graphpad Prism. The current-voltage relationships were plotted as normalized steady state current values at the end of the pulse step (at 400 ms) versus the respective voltage step (mV). Current densities (pA/pF) were obtained by dividing the steady state current values(pA) by the cell membrane capacitance (pF). The conductance voltage relationship was plotted to analyze the voltage dependence of channel activation. The conductance (G) was calculated using I=G(V_t_-V_rev_), where I is the steady state current value at the test potential V_t_, and V_rev_ is the reversal potential. The conductance voltage plot was fitted to a Boltzmann equation: I=I_max_/1+exp((V_t_-V_h_)/k)) where I_max_ is the maximum steady state current amplitude, V_h_ is half maximal voltage for activation(V_50_), and *k* is the slope factor. The number of recordings (n) is mentioned in the figure; a two-tailed unpaired t-test is used for calculating the significance, ***p < 0.001; n.s., not significant.

The conductance density was calculated by dividing the conductance values (nS) by the cell membrane capacitance (pF).

## Data Availability

The structures have been deposited with the accession numbers in PDB, 8GTR(PANX3), 8GTS(PANX1_R217H_), 8GTT(PANX1_DM_). The CryoEM density maps have been deposited in EMDB under accession numbers, EMD-34265(PANX3), EMD-34268(PANX1_WT_), EMD-34266(PANX1_R217H_), EMD-34267(PANX1_DM_).

Source data for the images is also uploaded along with the manuscript.

## Figure Legends Main Figures

## Supplementary Figures

**Figure S1:** Multiple sequence alignment of human PANX isoforms. The mutants studied are marked; PANX1_DM_, represents a double mutant (W74R, R75D). PANX1_R217H_ is a congenital mutant, PANX1_R24A_ PANX1_R29A_ are N-terminus mutants, PANX1_K36A_ is a TM1 mutant, PANX1_R75A_ is a pore mutant, and PANX1_R128A_ is a substitution at TM2. TM= Transmembrane, NTH= N-terminal Helix, ICH=Intracellular helix, CTH=C-terminal helix.

**Figure S2:** CryoEM processing workflow for PANX3; Initial ab-initio was done without symmetry. As seven subunits could be visualized in the ab-initio 3D reconstruction, C7 symmetry was applied for ab-initio and non-uniform refinement for the final reconstruction. A total of 31k particles were used for the final 3D reconstruction.

**Figure S3:** Representative densities for PANX3 at 7.5 σ. The secondary structures are labelled according to the topology diagram of PANX3 (**Figure S4A**).

**Figure S4:** Structural features of PANX isoforms **(A)**, Topology diagram of PANX1(blue), PANX2(green) and PANX3(orange), yellow dots represent the position of cysteines involved in disulfide bond formation in PANX3. TM= Transmembrane, NTH= N-terminal Helix, ICH=Intracellular helix, CTH=C-terminal helix. **(B)**, The positions of disulfide bonds in PANX3; SS1(66-261), SS2(84-242) **(C)**, Top view of superposed PANX1 and PANX3 displaying the differences in the position of N-glycosylation in PANX1(blue) and PANX3(orange). **(D-E)**, Density for NAG, and POPE contoured at 7.5 σ level.

**Figure S5**: Comparison of PANX3 with PANX1 and 2 isoforms. **(A)**, Superposition of PANX3(orange) and PANX3 Alphafold2 model(teal) reveals that N-terminus is towards the cytoplasmic side. **(B)**, Superposition of PANX3(orange) and PANX2 (brown) shows that R89 and D90 are the pore residues in PANX2. **(C)**, Multiple sequence alignment showing W74 is not conserved between PANX1,2 and PANX3 **(D)**, PANX3 displaying a gap between the protomers; the gap is large enough to accommodate ions and can act as a side tunnel; Inset shows the superposition of PANX1 and PANX3 along with PANX2 and PANX3, displaying the position of the side tunnel. **(E)**, Multiple sequence alignment exhibiting I58 is not conserved between PANX1, 2 and PANX3, **(F)** An annulus of seven uncharacterized densities in PANX3 contoured at 7 sigma.

**Figure S6:** Biochemical analysis of PANX1 mutants. **(A)** MST binding experiments for PANX1_WT_ with ATP, **(B-C)** Binding affinity with ATP-γs for PANX1 and PANX3 was determined as 13±3 and 75±19 μM, respectively, **(D)**, MST binding experiments for PANX1_WT_ with CBX (carbenoxolone), highest concentration was kept at 5mM, however no saturation was observed **(E)**, The position of the mutants selected for ATP-γS binding is displayed **(F)**, buffer was kept as a control (**G)**, Microscale thermophoresis profile for PANX1_R24A_ shows the binding of 83 ± 14μM. **(H)**, The binding affinity for the putative ATP binding site (R75) in PANX1 was determined to be 75± 19 μM, suggesting that R75 is not the sole residue responsible for ATP-γS binding in PANX1, **(I)**, The binding affinity for the PANX1 double mutant was determined to be 78± 29 μM **(J)**, Microscale thermophoresis profile for PANX1_R128A_ displays a complete loss of ATP-γS binding. **One of three independent experiments is shown in these figures. Repeats are displayed in Figure S14**.

**Figure S7:** Patch clamp studies of PANX1 and PANX3 **(A)**, Percentage inhibition by a PANX inhibitor, CBX plotted at +100mV for wild-type PANX1, 3 and the mutants along with untransfected controls. The number of recordings is mentioned in the graph; error bars represents SEM. A two-tailed unpaired t-test is used for calculating the significance, ***p < 0.001; n.s., not significant. (**B)**, Current density is plotted for the PANX1_WT_(n=8) and the mutants, PANX1_R75A_(n=8), PANX1_R217H_(n=6), PANX1_R24A_(n=4), PANX1_R128A_(n=6), PANX1_DM_(n=8), untransfected (n=6), the error bar represents SEM. **(C)**, Conductance density-Voltage plot for the PANX1_WT_ and the mutants is shown. Each point represents the mean of n =4-5 individual recordings, and the error bar represents SEM **(D)**, Normalized Conductance-voltage (GV-curve) plot for PANX1_WT_(n=8), PANX1_R75A_(n=6), and PANX1_DM_ (n=4) suggests that the pore residues are not involved in the voltage sensitivity of the channel, Conductance-voltage(GV-curve) plot for PANX1_R217H_ exhibits a reduction in the voltage sensitivity of the channel, n=4; the error bar represents SEM. **(E)**, The normalized G-V values were fitted with the Boltzmann equation, and the voltage at which the half-maximal activation, V_50_, occurred along with slope factor, *k*, was calculated for all the constructs.

**Figure S8**: Substitutions in PANX1 vestibule alter ATP interactions and channel behaviour **(A)**, Multiple sequence alignment of PANX1_WT_ displaying arginine conservation at 217 position. **(B)**, Fluorescence size exclusion chromatography (FSEC) profile for PANX1_WT_ (blue) on superose6 increase column, the heptamer, and the free GFP is marked on the profile. **(C)**, Fluorescence size exclusion chromatography (FSEC) profile for PANX1_WT_ and the mutants **(D)**, PANX1 double mutant through W74/R75 substitution, A cross-section of superposed PANX1_WT_ and PANX1_DM_ displays the reduction of pore; the density for W74R **(E)**, The structural superposition of the PANX1_WT_ and the mutant displays the changes in the movement leading to the reduced pore size; for clarity, only one subunit is shown. **(F)**, A cross-section of superposed PANX2 and PANX1_DM_ displays the position of arginine at the pore.

**Figure S9:** CryoEM workflow for the PANX1_WT_ and the mutants. **(A)**, The structure of PANX1_WT_ was determined at a resolution of 3.75 Å with C7 symmetry; a total of 41k particles were used for the 3D reconstruction. **(B)**, A similar workflow was used for the congenital mutant, PANX1_R217H_; 40K particles were used for the final 3D reconstruction for a final resolution of 3.87 Å. **(C)**, The PANX1 double mutant, PANX1_DM_ was determined at a resolution of 4.29 Å with a total of 41K particles. All the processing was done in Cryosparc.

**Figure S10:** Raw data of patch clamp studies performed with PANX1 and PANX3. **(A)**, The protocol used for patch clamp is shown; the voltage steps of 10/20mV were applied from - 120 to +120, **(B)**, mock recordings(untransfected HEK293 cells) were done prior to the PANX1_WT_ and mutant experiments; **(C-G)**, Representative raw traces along with the current density-voltage plot for whole-cell current for HEK cells expressing PANX1_WT_ and the mutants are displayed, 100μM CBX was applied to study the effect of the inhibitor on PANX channels.

**Figure S11:** Size exclusion profiles of PANX1 and PANX3 constructs. **(A)**, Size exclusion chromatography (SEC) profile for the PANX1_WT_ (blue), the fractions used for cryoEM grid preparation are marked. **(B)**, SEC profile for congenital mutant PANX1_R217H_ is similar to that of PANX1_WT_. **(C)**, SEC profile for PANX1_DM_, double mutant, the main fraction is taken for the grid freezing. **(D)**, SEC profile for PANX3, we observed heterogeneity in the protein, and the protein corresponding to PANX3, labeled as cryoEM, was used for grid freezing. **(E-G)**, SEC profiles for PANX1 mutants used for ATP binding studies, PANX1_R24A_(pink), PANX1_R128A_(gray), and PANX1_R75A_(green). All the proteins were run on a superose6 column.

**Figure S12: (A)**, FSEC profile for PANX isoforms (PANX1,2 and 3 solubilized in GDN), (**B)**, Structural comparison of Hs PANX1_WT_ (PDB: 6WBF(cyan) and and Zf PANX1_WT_,, 6VD7(orange)) (**C)**, Structural comparison between existing Hs PANX1_WT_. (PDB: 6WBF(cyan), 6V6D(pink), 7DWB(yellow).

**Figure S13**, SDS-PAGE profile for the PANX1_WT_, PANX3 and the PANX1 mutants used in the study.

**Figure S14**, MST analysis for the PANX1_WT_, PANX1 mutants and PANX3 with ATP-γS. The experiments are done in triplicates. The individual MST experiments with binding affinities are shown. A representative figure from the triplicates is shown in the Figure3 and the Figure S6.

**Movie S1**. A top down view of PANX1_R217H_ displaying rotameric shifts of W74 at the pore as a morph between PANX1_WT_ to PANX1_R217H_ that constricts the channel opening.

**Table S1**. CryoEM data collection and refinement statistics.

