## Supplementary figure 1-14; Table S1 for "Structural insights into the organization and channel properties of human Pannexin isoforms 1 and 3"

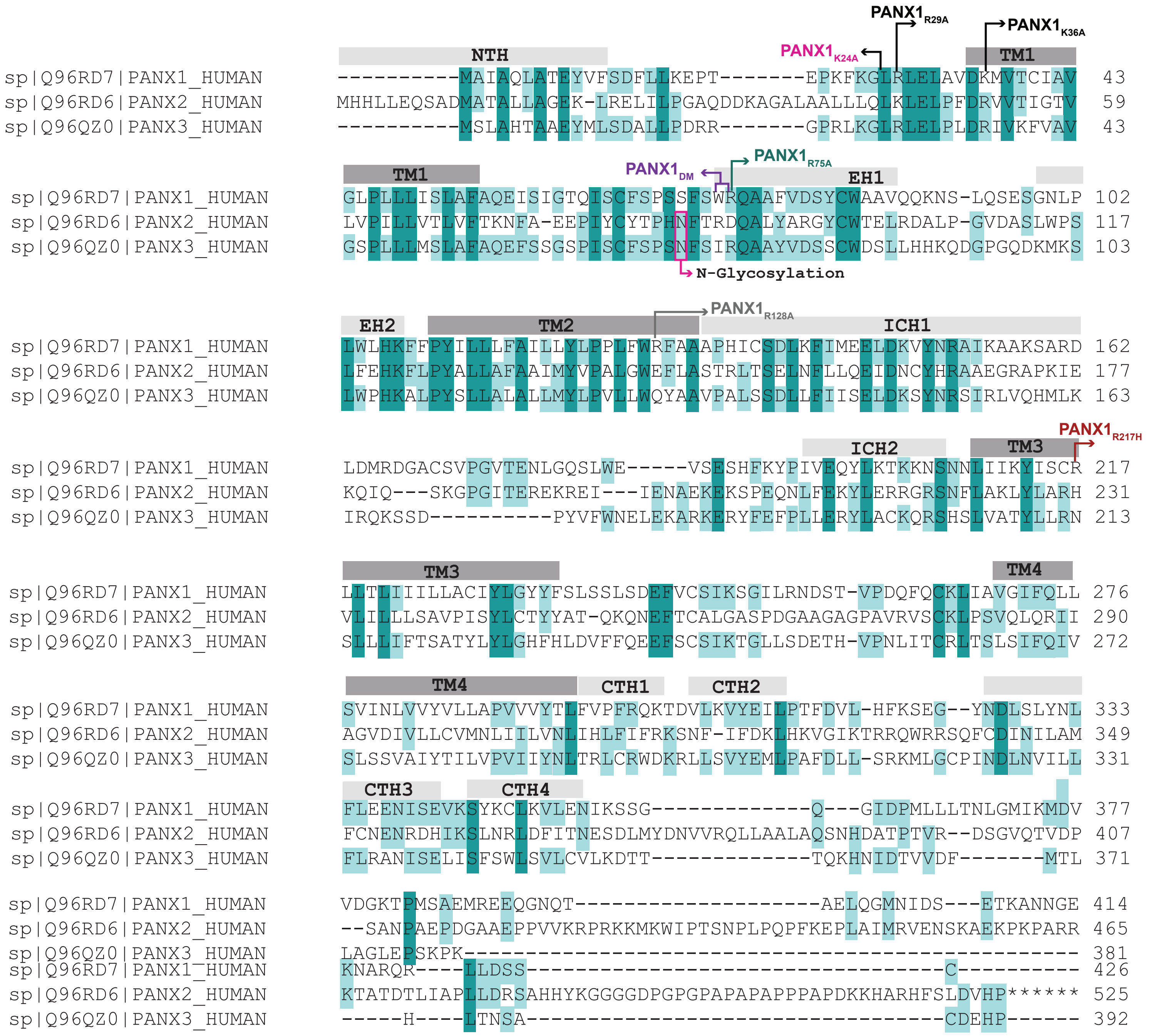


**Figure S1: Multiple sequence alignment for PANX isoforms(PANX1, 2 and 3)**

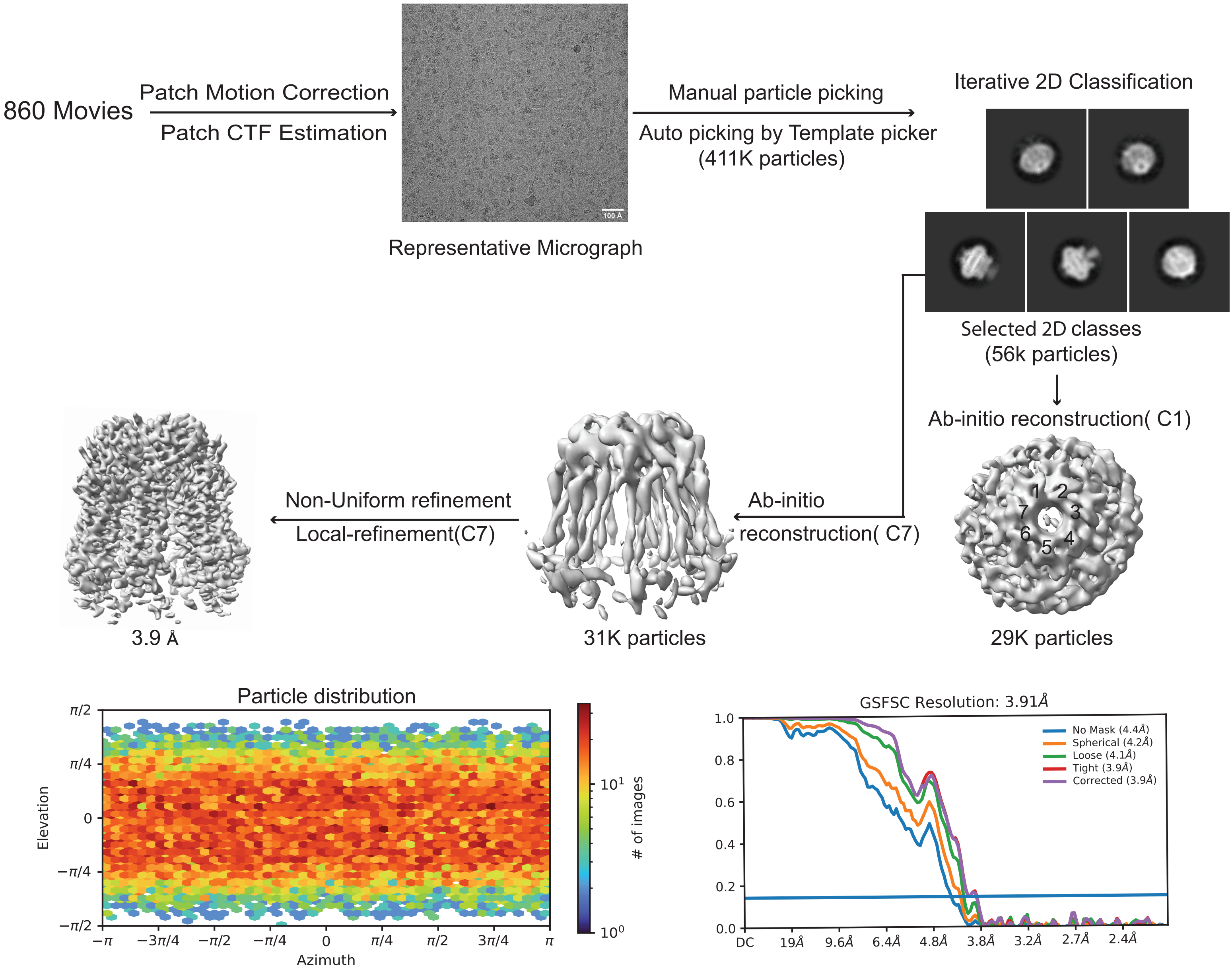


**Figure S2: Cryo-EM workflow for PANX3 data processing.**

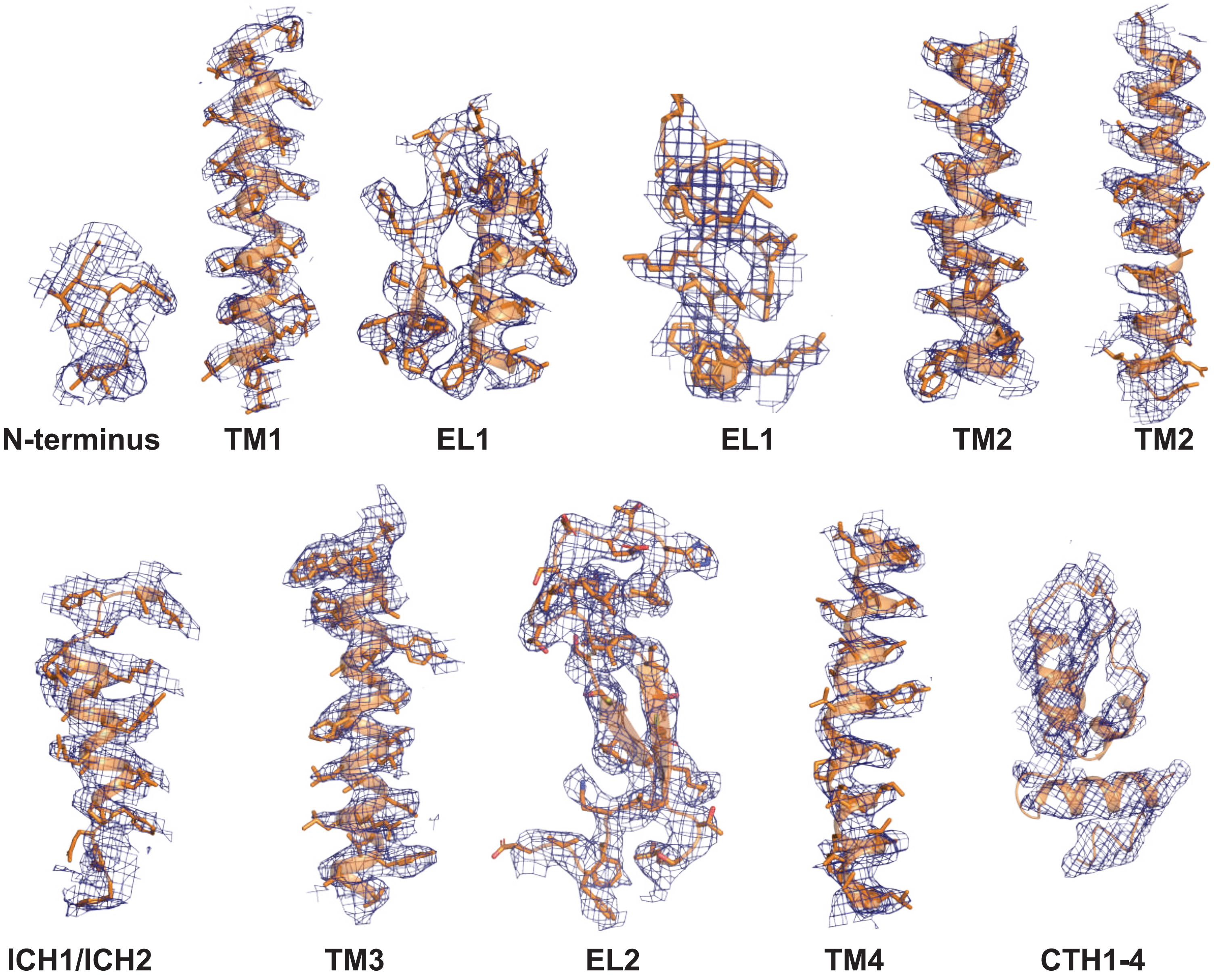


**Figure S3: Representative densities for PANX3.**

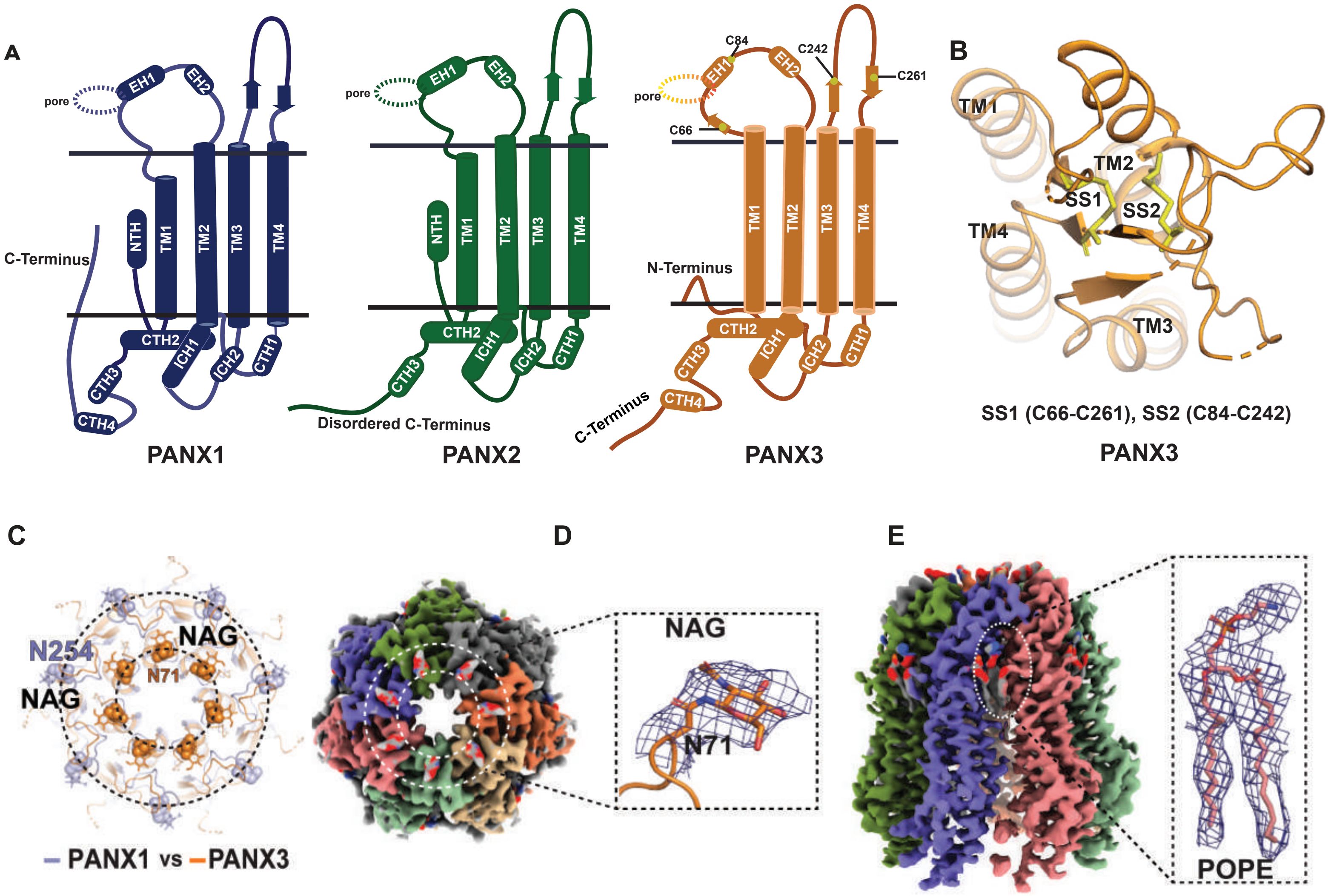


**Figure S4: Structural features of PANX1, 2 and PANX3**

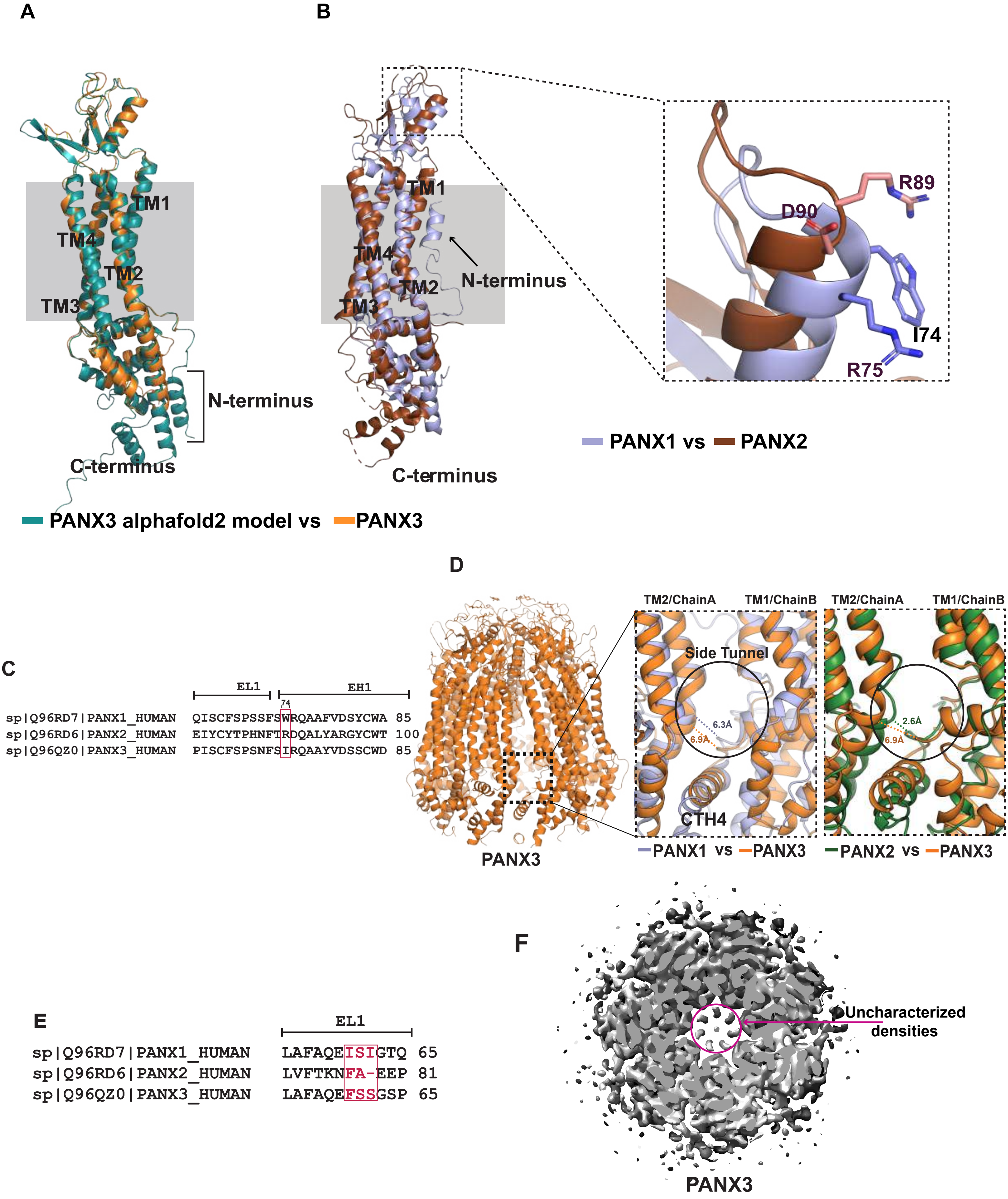


**Figure S5: Comparison of PANX3 with PANX1 and PANX2 isoforms**

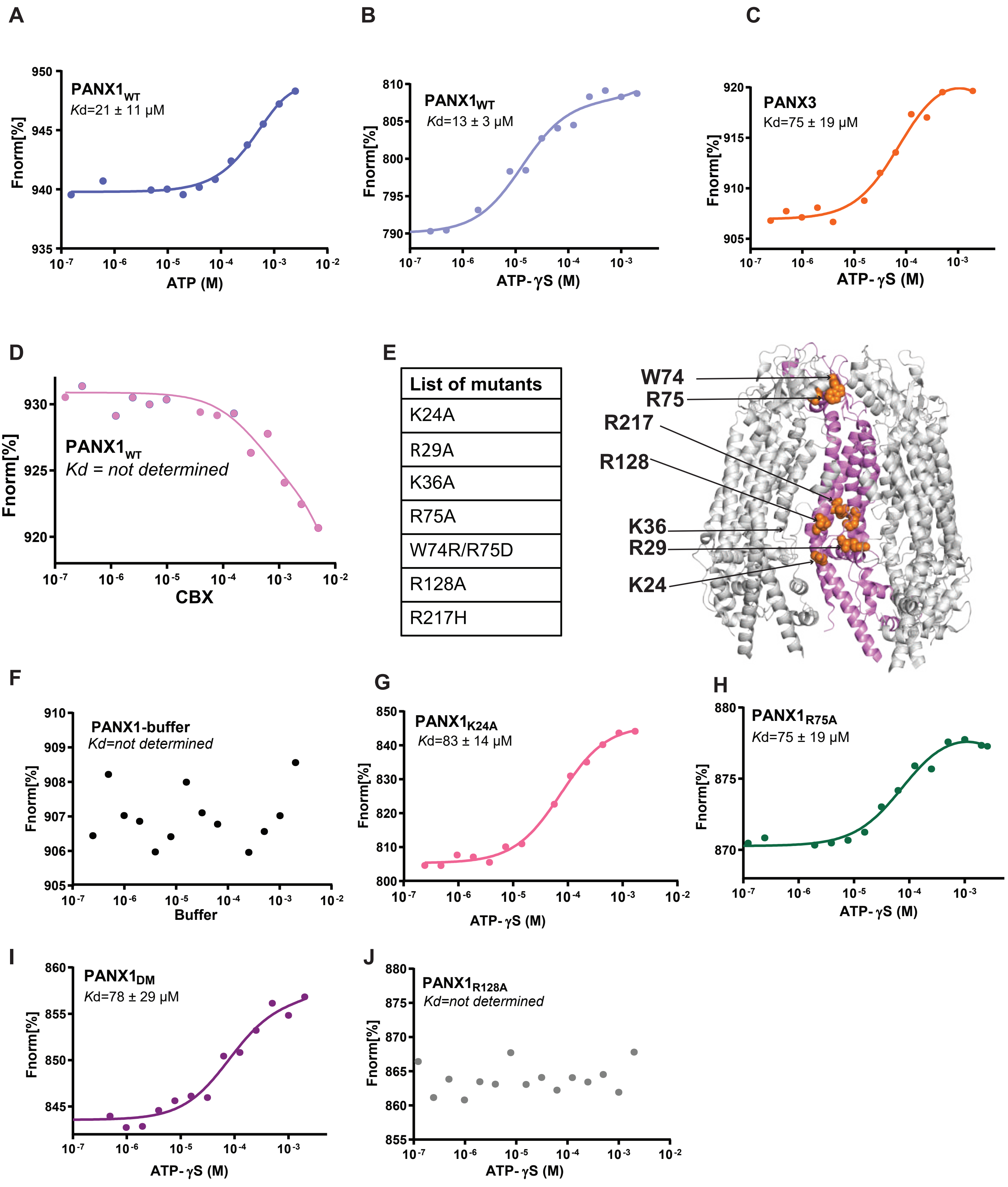


**Figure S6: Biochemical analysis of PANX1 and 3 isoforms and PANX1 mutants.**

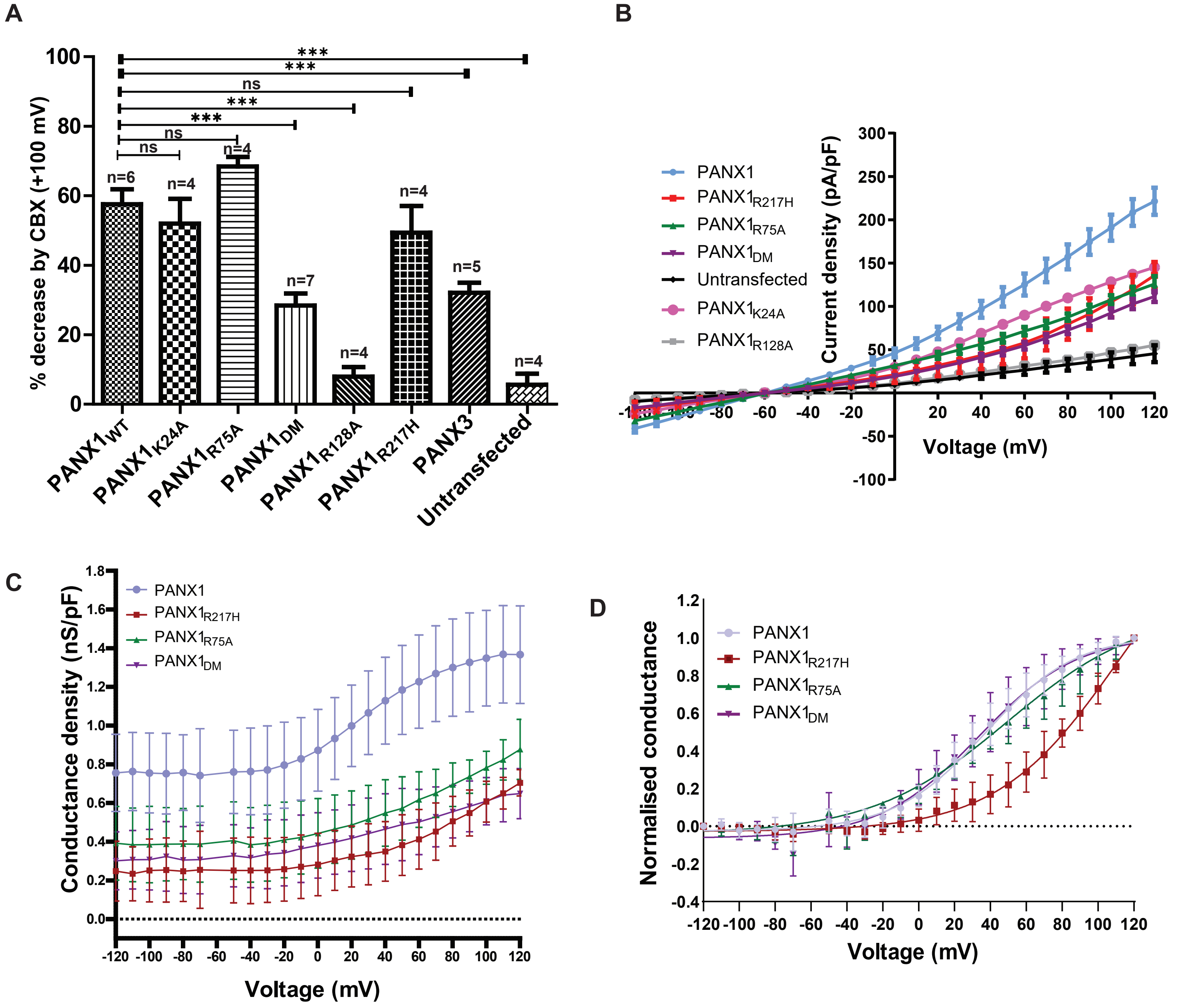


E

| Constructs | $V_{50}$ | Slope factor, k |
|---|---|---|
| PANX1$_{WT}$ | 36.92 ± 1.89 | 26.03 ± 1.7 |
| PANX1$_{R75A}$ | 49.7 ± 6.72 | 38.17 ± 5.5 |
| PANX1$_{DM}$ | 32.34 ± 3.96 | 29.47 ± 3.8 |
| PANX1$_{R217H}$ | 115.9 ± 18.55 | 35.63 ± 5.2 |

**Figure S7: Patch clamp studies for the PANX1 and PANX3**

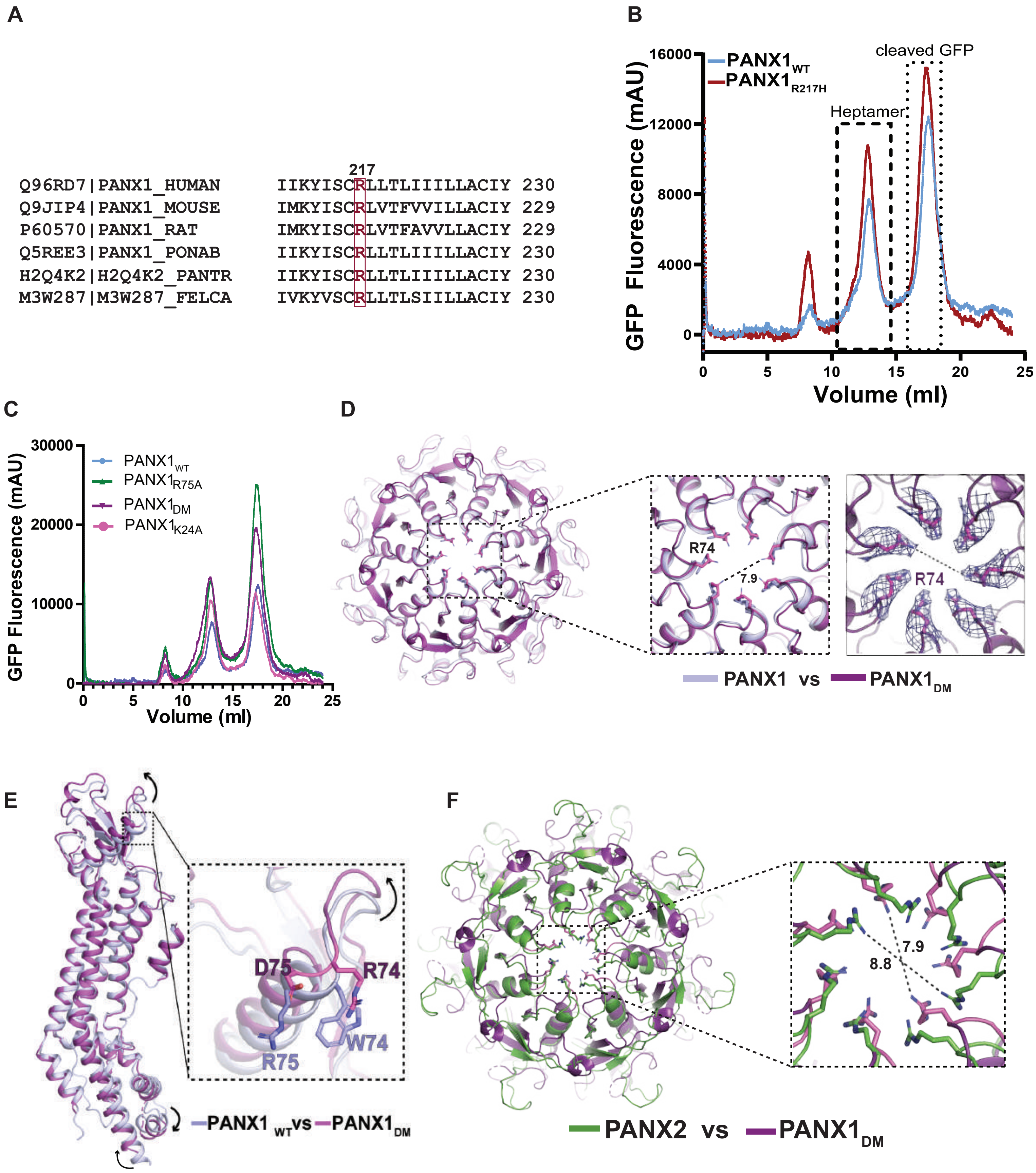


**Figure S8:Substitutions in PANX1 vestibule alter ATP interactions and channel properties.**

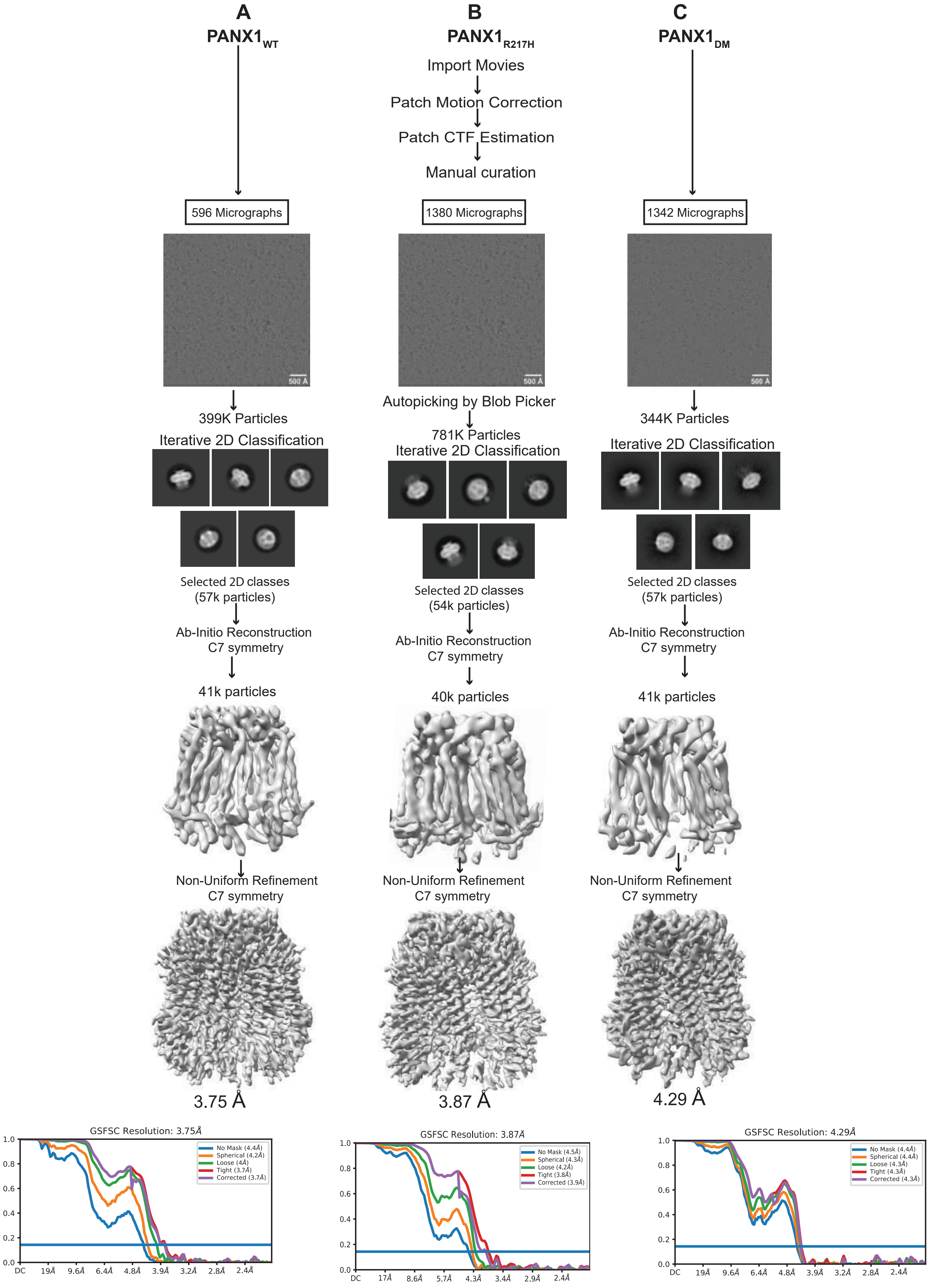


**Figure S9: Cryo-EM workflow for $PANX1_{WT}$ and mutants**

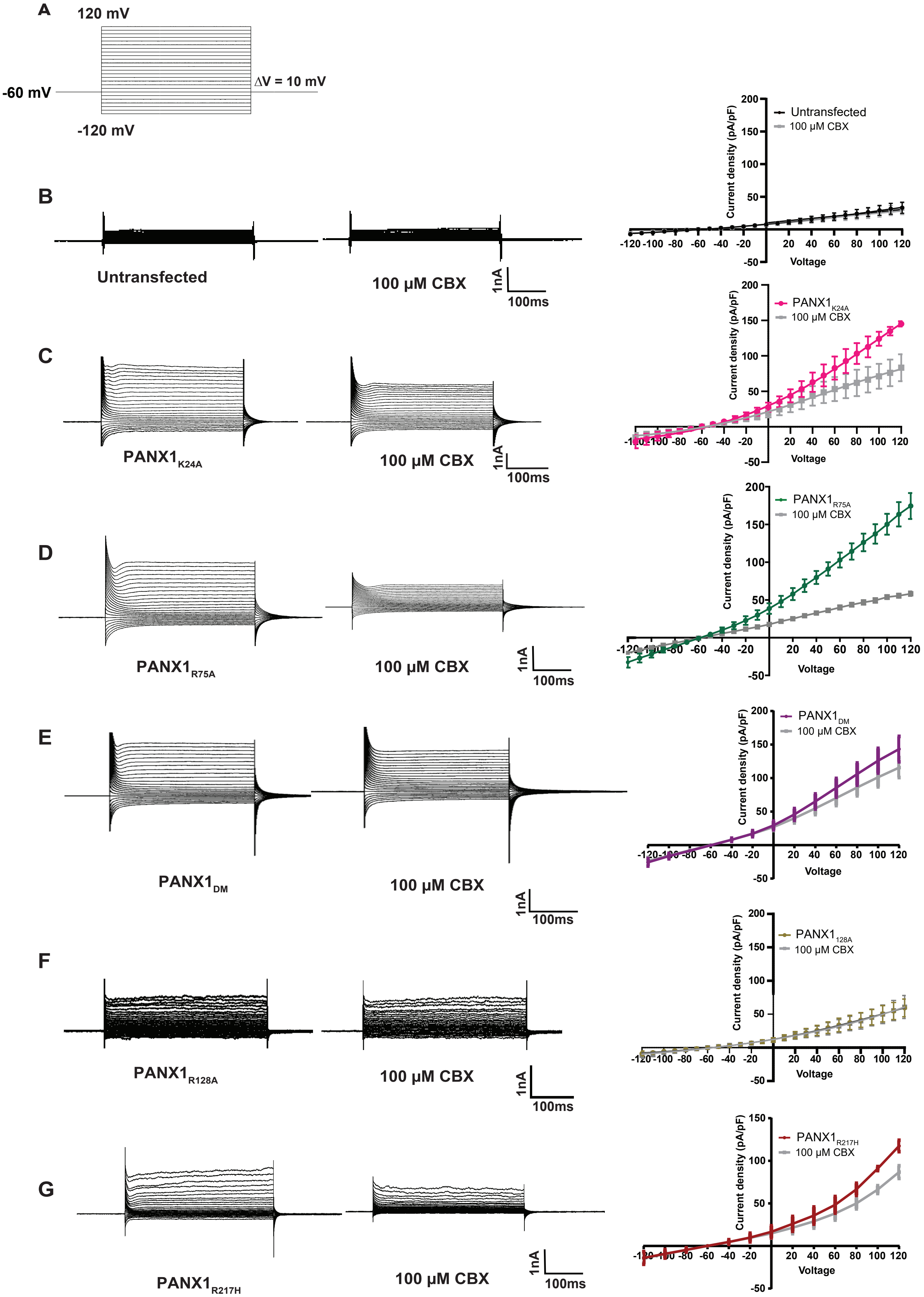


**Figure S10: Raw data of Patch clamp studies of PANX1 mutants.**

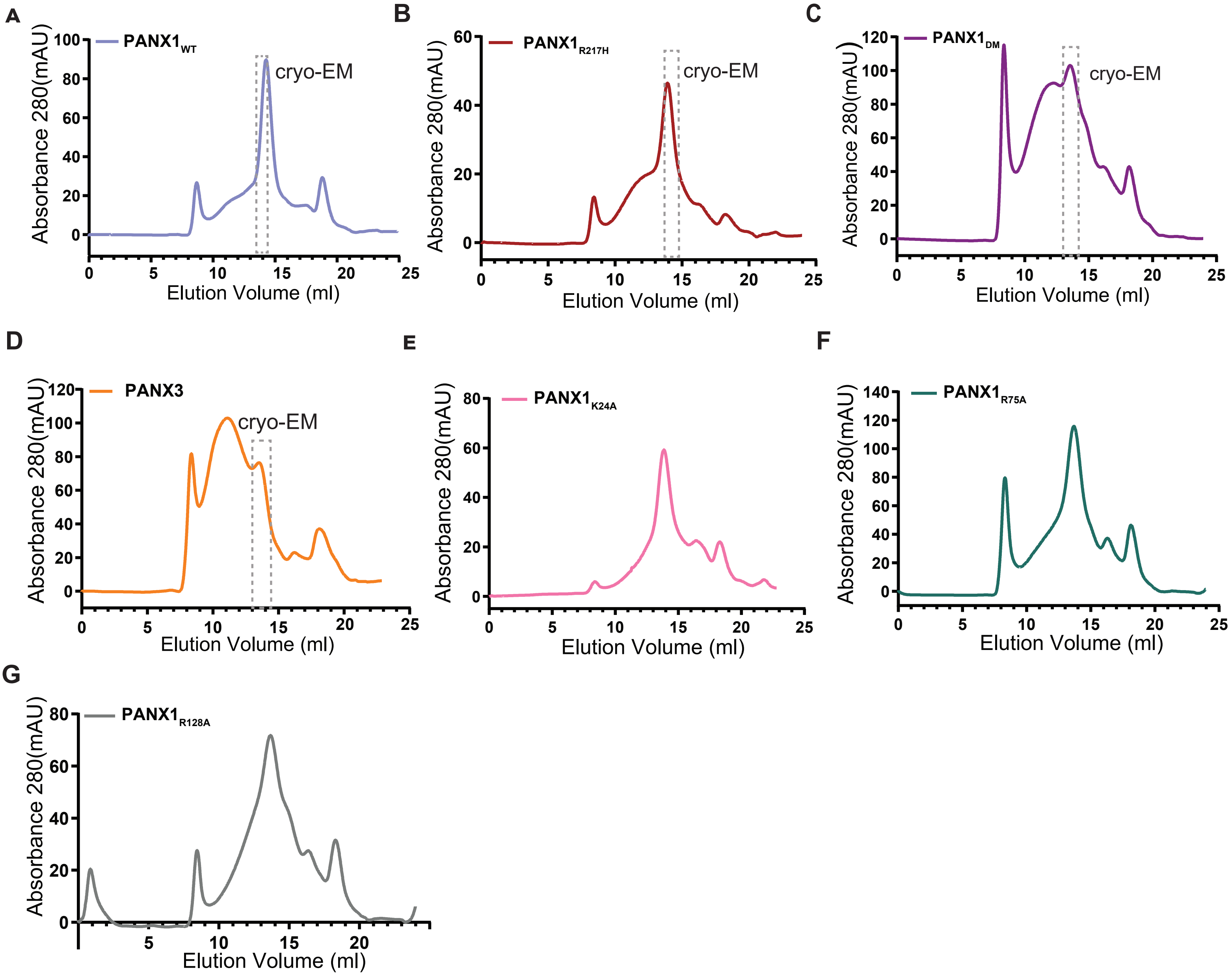


**Figure S11: Size exclusion chromatography profile for PANX1 and PANX3 constructs.**

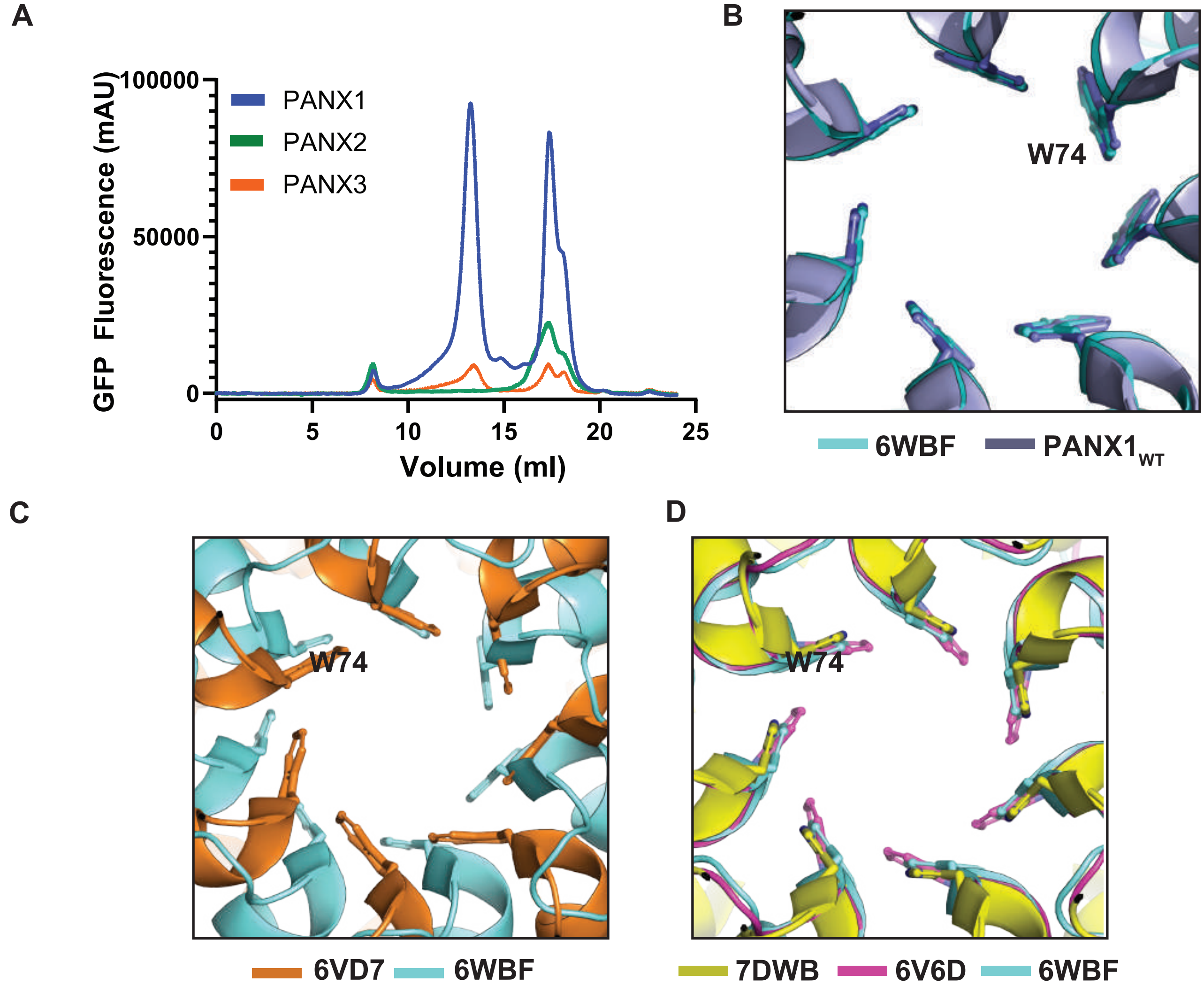


**Figure S12: Structural comparison of PANX channels.**

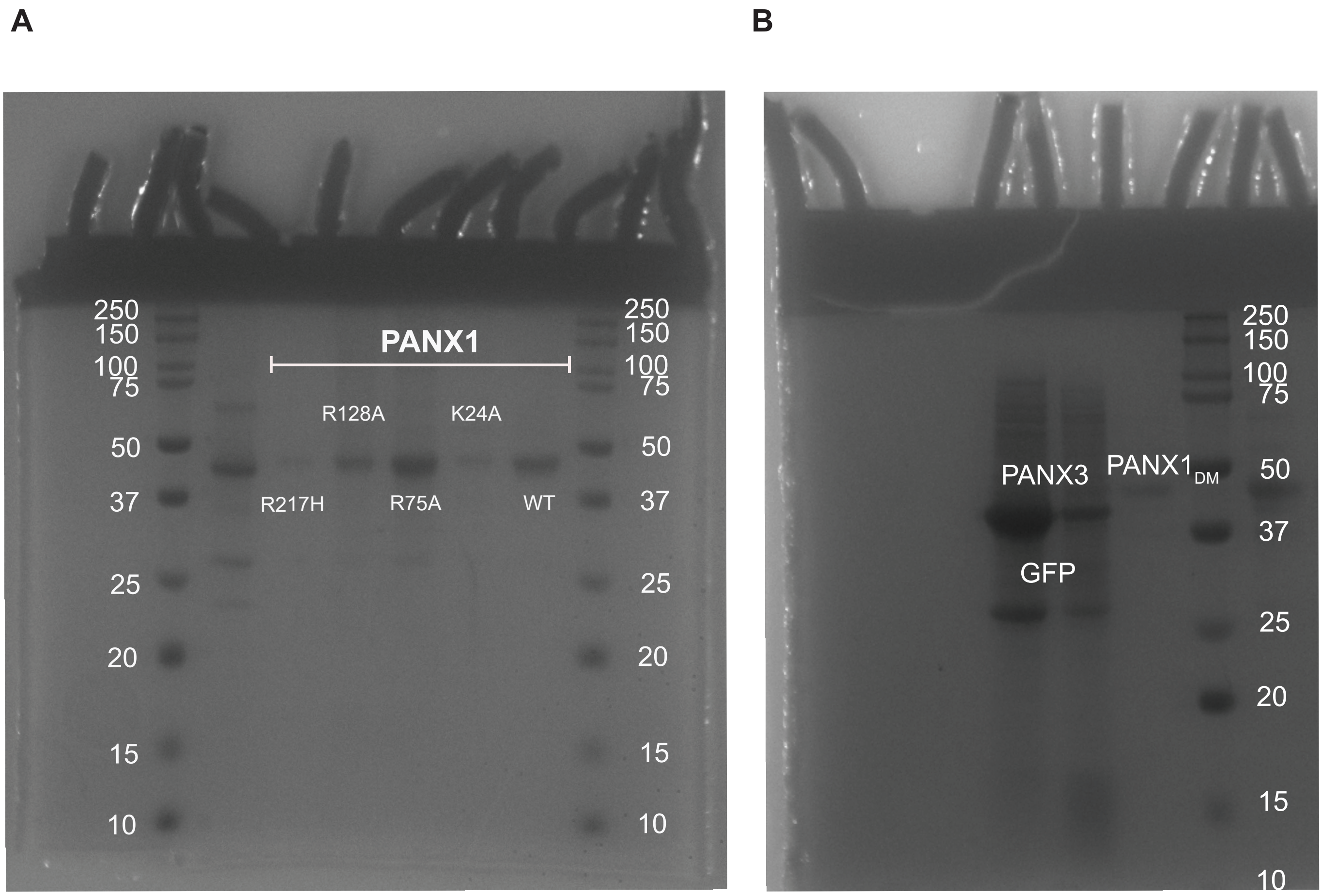


**Figure S13: SDS-PAGE profile for the constructs mentioned in the study.**

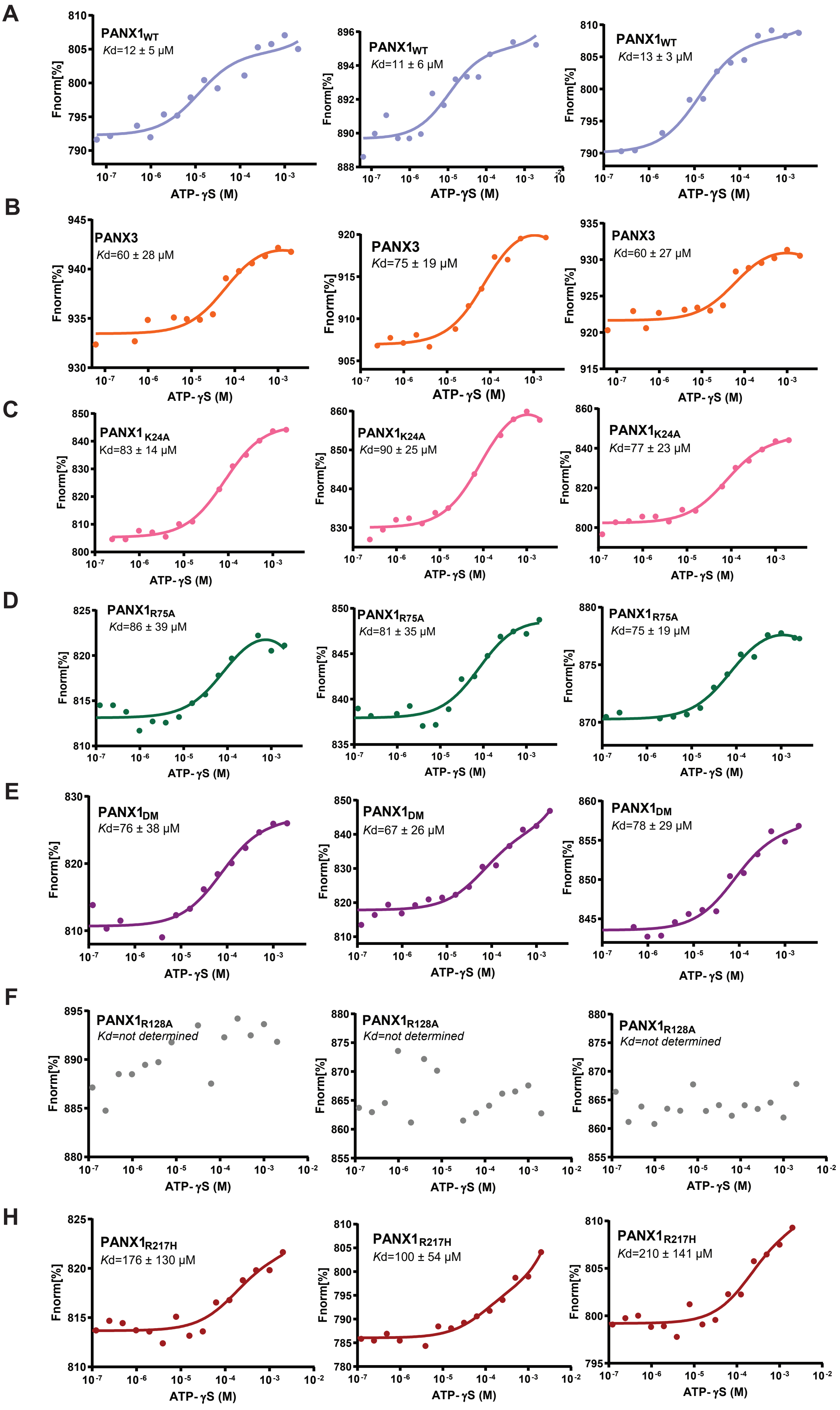


**Figure S14: MST analysis for the constructs used in the study.**

### Cryo-EM data collection, refinement and validation statistics

| | $PANX1_{WT}$ (EMD-34268) | $PANX1_{DM}$ (EMD-34267) (PDB 8GTT) | $PANX1_{R217H}$ (EMDB-34266) (PDB 8GTS) | PANX3 (EMDB-34265) (PDB 8GTR) |
|---|---|---|---|---|
| **Data collection and processing** | | | | |
| Magnification | 75000 | 75000 | 130000 | 130000 |
| Mode | TEM | TEM | EFTEM | EFTEM |
| Voltage (kV) | 300 | 300 | 300 | 300 |
| Detector | FalconIII | FalconIII | K2 | K2 |
| Slit width (eV) | - | - | 20 | 20 |
| Electron exposure ($e^{-}/Å^2$) | 29.5 | 29.87 | 41.12 | 48.8 |
| Defocus range (μm) | -1.8 to -3.3 | -1.8 to -3.3 | -1.8 to -3.3 | -1.8 to -3.3 |
| Pixel size (Å) | 1.07 | 1.07 | 1.07 | 1.07 |
| Micrographs (No.) | 1201 | 1404 | 1776 | 1112 |
| Symmetry imposed | C7 | C7 | C7 | C7 |
| Initial particle images (no.) | 399706 | 344639 | 781859 | 411263 |
| Final particle images (no.) | 41483 | 41282 | 40873 | 31517 |
| Map resolution (Å) | 3.75 | 4.29 | 3.87 | 3.91 |
| FSC threshold | 0.143 | 0.143 | 0.143 | 0.143 |
| **Refinement** | | | | |
| Initial model used (PDB code) | | 6WBF | 6WBF | Alphafold2 |
| Model resolution (Å) @ FSC 0.5 | | 4.4 | 4.2 | 4.2 |
| Map sharpening *B* factor ($Å^2$) | -179.6 | -141.1 | -156.0 | -143.6 |
| Model composition | | | | |
| Non-hydrogen atoms | | 16212 | 15470 | 16975 |
| Protein residues | | 2156 | 2016 | 2156 |
| Ligands | | - | - | 7 (NAG) 7 (PTY) |
| B-factor ($Å^2$) | | | | |
| Total | | 88.11 | 102.2 | 70.00 |
| Protein | | 88.11 | 102.2 | 70.57 |
| Ligands | | - | - | 42.92 (NAG) 46.01 (PTY) |
| R.m.s. deviations | | | | |
| Bond lengths (Å) | | 0.004 | 0.004 | 0.004 |
| Bond angles (°) | | 1.049 | 1.057 | 1.011 |
| Validation | | | | |
| MolProbity score | | 1.76 | 1.65 | 1.75 |
| Clashscore | | 10.64 | 10.73 | 7.72 |
| Poor rotamers (%) | | 0 | 0 | 0 |
| Ramachandran plot | | | | |
| Favored (%) | | 96.62 | 97.48 | 95.33 |
| Allowed (%) | | 3.38 | 2.52 | 4.67 |
| Disallowed (%) | | 0 | 0 | 0 |
